# Improving polygenic risk prediction performance through integrating electronic health records by phenotype embedding

**DOI:** 10.1101/2025.08.05.668705

**Authors:** Leqi Xu, Wangjie Zheng, Jiaqi Hu, Yingxin Lin, Jia Zhao, Gefei Wang, Tianyu Liu, Hongyu Zhao

## Abstract

Large-scale biobanks provide comprehensive electronic health records (EHRs) that capture detailed clinical phenotypes, potentially enhancing disease risk prediction. However, traditional polygenic risk score (PRS) methods rely on simplified phenotype definitions or predefined trait sets, limiting their ability to represent the intricate phenotypic structures embedded within EHRs. To address this gap, we introduce a general framework, EEPRS, that leverages phenotype embeddings derived from EHRs to improve genetic risk prediction using only genome-wide association study (GWAS) summary statistics, enabling accurate, robust and interpretable risk prediction for a wide range of diseases. Employing embedding methods such as Word2Vec and GPT, we conducted EHR embedding-based GWAS and identified a distinct cardiovascular cluster via hierarchical clustering of genetic correlations. Across 41 clinical traits in the UK Biobank, our EEPRS framework consistently outperformed traditional single-trait PRS, particularly within this identified cluster. Validation using PRS-based phenome-wide association studies (PRS-PheWAS) further confirmed robust associations between EHR embedding-based PRS and circulatory system diseases. Furthermore, our data-adaptive method, EEPRS_optimal, employing cross-validation to select the best embedding method, leading to additional improvements in prediction. We further developed MTAG_EEPRS for multi-trait PRS, resulting in averaging 92.48% improvement in *R*^2^ for continuous traits and 24.06% in AUC for binary traits compared to single-trait PRS. Overall, EEPRS represents a robust and interpretable framework, enhancing genetic prediction accuracy through integrating EHR embeddings with single-trait and multi-trait PRS.

## Introduction

Large-scale biobanks, such as the UK Biobank (UKBB) [1] and the Million Veteran Program (MVP) [2], provide comprehensive electronic health records (EHRs) comprising structured data (e.g., diagnosis codes) and unstructured data (e.g., clinical notes). Integrating these rich clinical profiles with genetic data presents a great opportunity to accelerate genetic discoveries, enhance disease risk prediction, and improve personalized treatment [3].

Despite this potential, current genome-wide association studies (GWAS) often define phenotypes based on simplified criteria, typically grouping related ICD-10 diagnosis codes into binary disease definitions (case vs. control) [4]. This simplification potentially neglects the intricate phenotypic relationships embedded in detailed patient records, limiting both the statistical power and clinical utility of genetic studies.

This limitation is particularly evident in traditional polygenic risk scores (PRS), which aggregate effects across multiple genetic variants based on GWAS summary statistics typically derived from single diseases or traits [5, 6, 7, 8, 9]. As a result, these single-trait PRS methods may overlook the pleiotropic effects and shared genetic signals across multiple phenotypes available within EHRs [10, 11]. While existing multi-trait PRS methods partially address these limitations, they remain constrained by predefined phenotype sets, unable to fully exploit the comprehensive and high-dimensional clinical data in EHRs [12, 13, 14, 15, 16].

For instance, coronary artery disease (CAD) risk prediction commonly incorporates PRS derived from CAD, related atherosclerotic diseases (e.g., ischemic stroke), and established risk factors such as diabetes, blood pressure, and lipid concentrations [17]. However, these preselected phenotypes may inadequately represent the broader clinical traits and their complex interrelations inherently captured by EHRs, thereby limiting PRS performance.

Recent studies have explored incorporating EHRs into genetic risk prediction frameworks [18]. However, they focus on single-trait summaries, such as spirograms, and neglect complex cross-trait phenotypic patterns within EHRs. Moreover, current methods typically require individual-level data for PRS integration, limiting their utility when only summary statistics are available [18]. Thus, methods that effectively leverage comprehensive clinical information from EHRs using summary statistics for genetic risk prediction remains largely unexplored.

With the advancements in embedding approaches, our ability to represent complex EHRs has been significantly enhanced [19]. Traditional natural language processing (NLP) methods, such as Word2Vec [20], and transformer-based large language models (LLMs), such as GPT-o1 [21, 22], can effectively capture semantic relationships within structured and unstructured EHRs, even in scenarios with incomplete information. By transforming clinical records into numeric vector embeddings, these methods encapsulate rich contextual and phenotypic relationships. Compared to binary ICD-10 disease indicators, these embeddings leverage the high-dimensional, heterogeneous nature of EHRs, capturing richer clinical contexts and improving the potential for genetic discovery and risk prediction.

Motivated by these developments, we propose the Electronic Health Record Embedding Enhanced Polygenic Risk Score (EEPRS) framework, a unified, interpretable, and flexible framework leveraging high-dimensional EHR information to enhance genetic risk prediction for both single and multiple traits. EEPRS introduces several innovations: (1) it transforms complex EHRs into lower-dimensional embeddings, enabling GWAS analyses directly on these embeddings to uncover phenotypic insights missed by single-trait GWAS; (2) it performs hierarchical clustering of clinical traits based on their genetic correlations with EHR embedding-based GWAS, identifying clinically and genetically coherent trait clusters; (3) it constructs and interprets EHR embedding-based PRS using PRS-based phenome-wide association studies (PRS-PheWAS) [23, 24]; (4) it combines EHR embedding-based PRS with traditional PRS using only GWAS summary statistics, eliminating the need for individual-level tuning data; (5) it utilizes a data-adaptive cross-validation strategy to systematically identify the optimal embedding method; and (6) it integrates flexibly with existing multi-trait PRS methods.

Our analyses demonstrate that EHR embedding-based GWAS detect significant heritability and genetic correlations with diverse clinical traits. Hierarchical clustering based on genetic correlation profiles between clinical traits and EHR embedding-based GWAS revealed a distinct cardiovascular cluster, capturing clinically and genetically coherent phenotypic architectures. By integrating EHR embedding-based PRS with traditional single-trait PRS within the EEPRS framework, we observed consistent improvement of prediction accuracy over single-trait PRS across all five embedding methods evaluated for 41 clinical traits in the UKBB European cohort [1]. PRS-based PheWAS further validated enhancement from EEPRS within the cardiovascular cluster, highlighting robust associations between EHR embedding-based PRS and circulatory system diseases. Additionally, our results underscore the advantage of the proposed data-adaptive embedding selection procedure and demonstrate the added predictive benefit of integrating multitrait PRS within the EEPRS framework.

## Results

### Overview of EEPRS

The EEPRS framework integrates EHR embeddings with single-trait and multi-trait PRS through five steps (Figure 1): (1) Generate EHR-based phenotype embeddings; (2) Perform EHR embedding-based GWAS; (3) Derive EHR embedding-based PRS; (4) Interpret EHR embedding-based PRS in a PRS-based PheWAS framework; and (5) Integrate EHR embedding-based PRS via EEPRS-Integrator in the EEPRS framework. Detailed implementations of EEPRS are provided in the Methods section.

**Figure 1:**
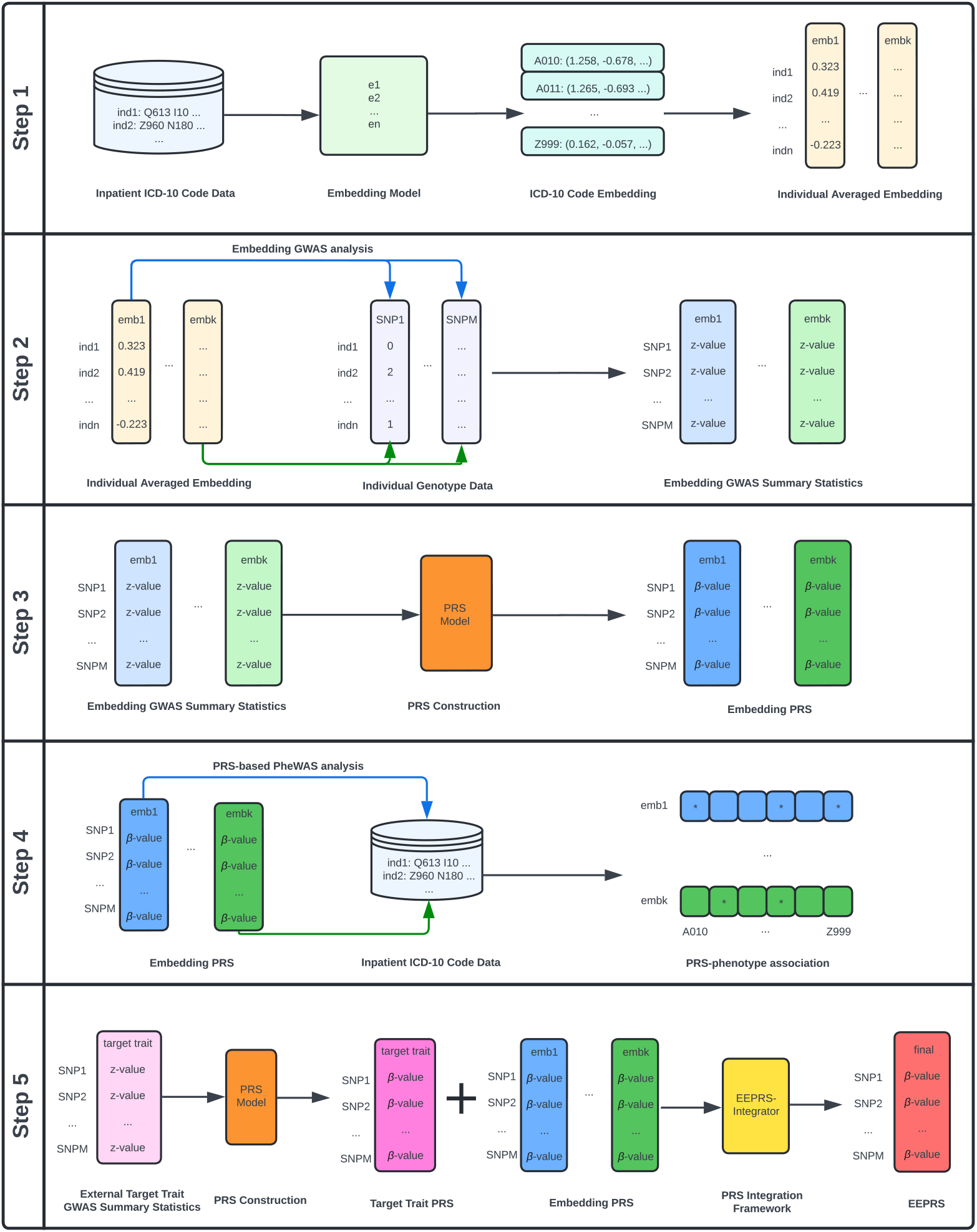
Overview of the EEPRS framework. EEPRS integrates EHR embeddings through a structured five-step workflow: (1) Generation of individual-level EHR embeddings from inpatient ICD-10 data; (2) Construction of EHR Embedding GWAS to identify SNP associations with embedding dimensions; (3) Construction of EHR embedding-based PRS using embedding GWAS summary statistics; (4) PRS-based PheWAS analysis to interpret EHR embedding-based PRS associations with inpatient ICD-10 data; (5) Integration of EHR embedding-based PRS and external target trait PRS using EEPRS-Integrator to produce final EEPRS estimates. This figure was created with Lucid.

#### Step 1: Generate EHR-based phenotype embeddings

To capture latent clinical associations and phenotypic co-occurrence patterns, we construct low-dimensional phenotype embeddings from EHRs. For each individual, these embeddings are derived from clinical descriptions associated with ICD-10 codes recorded in the patient’s diagnostic history, using embedding models such as Word2Vec [20] and GPT [21].

#### Step 2: Perform EHR embedding–based GWAS

To identify genetic signals potentially missed by conventional single-trait GWAS, we treat each embedding dimension generated in Step 1 as a quantitative phenotype and conduct GWAS analyses. To further identify clinically and genetically coherent trait clusters, we perform hierarchical clustering of clinical traits based on their genetic correlations with EHR embedding-based GWAS.

#### Step 3: Derive EHR embedding–based PRS

To quantify genetic predispositions associated with the phenotypic patterns captured by phenotypic embeddings, we derive EHR embedding-based PRS from the GWAS summary statistics generated in Step 2. Established PRS methods, such as PRS-CS-auto and SDPR [6, 8], can be applied, resulting in individual-level PRS vectors corresponding to each embedding dimension.

#### Step 4: Interpret EHR embedding-based PRS using a PRS-based PheWAS framework

To interpret the clinical and biological relevance of EHR embedding-based genetic risks, we perform PRS-based PheWAS analyses, evaluating associations between each EHR embedding-based PRS (predictor) and individual ICD-10 phenotypes (outcomes) [23, 24].

#### Step 5: Integrate EHR embedding-based PRS via EEPRS-Integrator in the EEPRS framework

As illustrated in Supplementary Figure 1, we integrate EHR embedding-based PRS with conventional PRS using EEPRS-Integrator. EEPRS-Integrator extends MIXPRS [25], a summary-level PRS integration method, by estimating optimal weights to combine EHR embedding-based PRS with traditional PRS without requiring individual-level tuning data. Additionally, to further enhance EEPRS performance, a data-adaptive approach employing cross-validation can be utilized to identify the optimal embedding strategy for each trait. Moreover, EEPRS is also flexible to incorporate multi-trait PRS results.

### EHR embeddings are heritable and genetically correlated with diverse traits

We first examined whether the phenotypic patterns represented by EHR embeddings capture heritable information and exhibit genetic correlations with complex traits, thereby offering potential to improve genetic risk prediction. We considered six embedding approaches to generate phenotypic representations from EHRs: (1) Word2Vec continuous bag-of-words (CBOW; Word2Vec) [20]; (2) Word2Vec CBOW reduced via principal component analysis (PCA; Word2Vec_PCA; capturing 80% variance) [26]; (3) Word2Vec CBOW reduced via independent component analysis (ICA; Word2Vec ICA; capturing 80% variance) [27]; (4) GPT embeddings derived by averaging word embeddings from GPT-o1-generated ICD-10 descriptions using text-embedding-3-large (GPT) [21, 22]; (5) GPT embeddings reduced via PCA (GPT_PCA; capturing 80% variance); and (6) GPT embeddings reduced via ICA (GPT_ICA; capturing 80% variance).

The UKBB dataset [1] was partitioned into training (two-thirds of European participants, *N* = 207, 734) and testing sets (one-third, *N* = 103, 867). EHR embeddings were generated from longitudinal clinical descriptions mapped to ICD-10 diagnosis codes using the training set, with embedding dimensions specific to each method: 100 dimensions for Word2Vec, 30 dimensions for Word2Vec_CA, 30 dimensions for Word2Vec_ICA, 3072 dimensions for GPT, 67 dimensions for GPT_PCA, and 67 dimensions for GPT_ICA. Unreduced GPT embeddings were excluded from subsequent analyses, as dimensionality reduction significantly reduced GPT embeddings from 3072 to 67 dimensions, indicating substantial redundancy and limited additional information in the unreduced embeddings.

We then performed GWAS for each embedding and estimated their heritability using linkage disequilibrium score regression (LDSC) [28]. As illustrated in Supplementary Figure 2, most embedding dimensions demonstrated statistically significant heritability, suggesting that the derived embeddings are genetically informative. Notably, ICA-derived embeddings contained dimensions exhibiting the highest heritability, specifically Word2Vec ICA (0.0570, SE = 0.0045) and GPT ICA (0.0638, SE = 0.0044).

Additionally, we evaluated genetic correlations between heritable embeddings and 51 clinical traits from diverse consortia, selected to represent major disease domains, biomarkers, and behavioral factors with broad relevance to human health [29, 30, 31, 32, 33, 34, 35, 36, 37, 38, 39, 40, 41, 42, 43, 44, 45, 46, 47, 48, 49, 50, 51, 52, 53, 54, 55, 56, 57, 58, 59, 60, 61, 62, 63] (Supplementary Table 1), using LDSC [64]. As shown in Figure 2, Supplementary Figures 3 and 4, most clinical traits exhibited significant genetic correlations with various EHR embedding-based phenotypes, underscoring the genetic relevance and potential clinical utility of these EHR embedding-based phenotypes.

**Figure 2:**
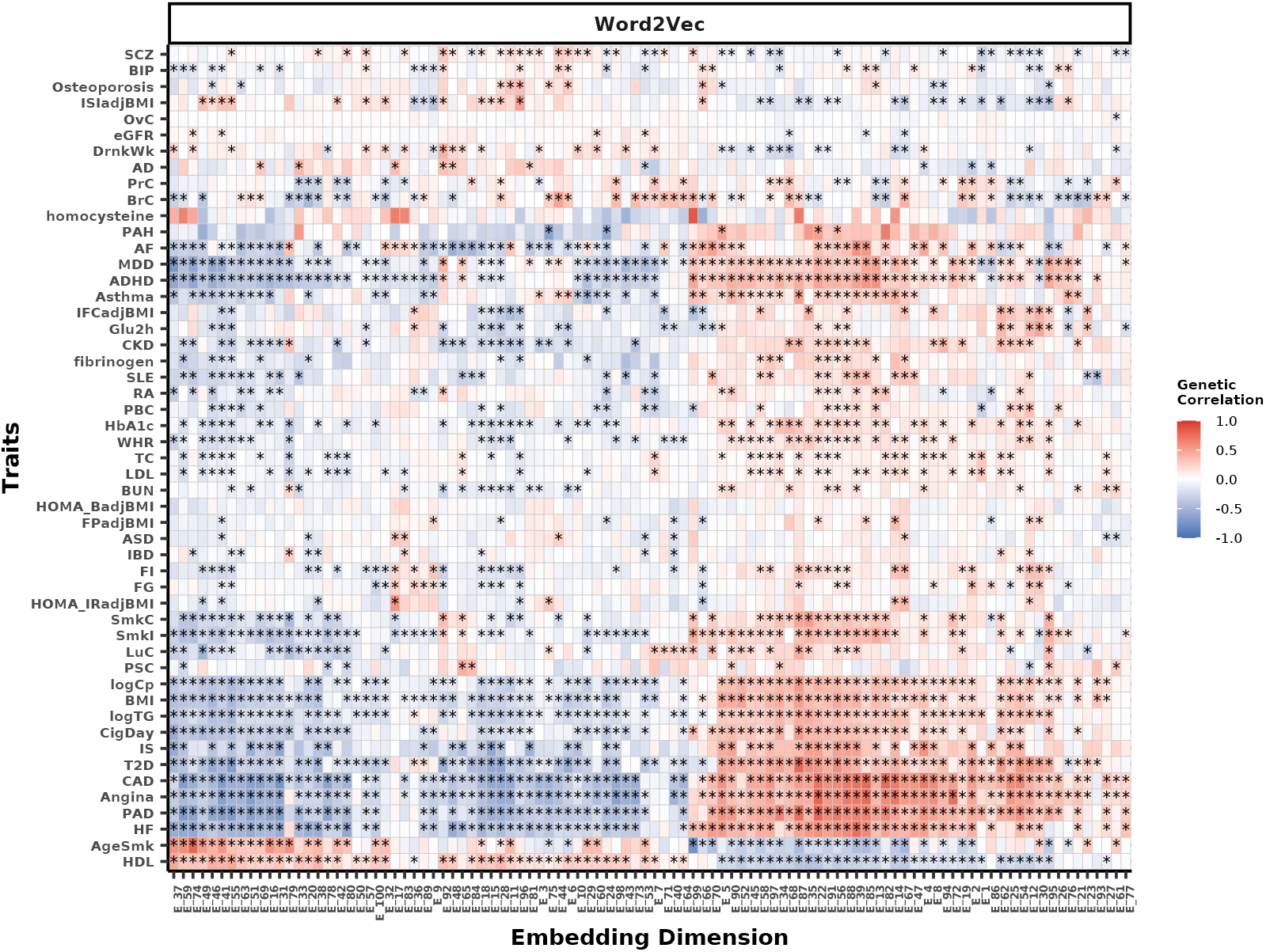
Genetic correlations between heritable Word2Vec embeddings and 51 complex traits. Heatmap illustrating genetic correlations between GWAS of heritable Word2Vec embeddings and 51 complex traits. Positive correlations are indicated in shades of red, and negative correlations in shades of blue, with color intensity reflecting correlation strength. Significant correlations (BH-adjusted p-value *<* 0.05) are marked by stars (*∗*).

To assess whether EHR embedding-based GWAS capture clinically and genetically coherent phenotypic structures, we performed hierarchical clustering of 51 traits based on their genetic correlation profiles with EHR-derived embedding dimensions. Consistently across five embedding methods, we identified six distinct trait clusters (Supplementary Figure 5). A particularly robust cluster, consistently observed in at least four embedding methods, grouped several canonical cardiovascular traits: coronary artery disease (CAD), ischemic stroke (IS), peripheral artery disease (PAD), angina, heart failure (HF), type 2 diabetes (T2D), body mass index (BMI), and log-transformed C-reactive protein (logCp). These traits share established clinical and biological connections, with conditions like T2D, elevated BMI, and increased CRP serving as well-recognized risk factors or markers for cardiovascular disease. Additionally, GWAS have revealed substantial polygenic overlap between T2D, BMI, and cardiovascular outcomes such as CAD [32, 54].

Together, these results highlight that genetic analyses based on embedding-derived GWAS effectively uncover known and meaningful genetic relationships across clinical traits. This approach demonstrates the potential for EHR-based embeddings to enrich genetic risk prediction through a scalable, data-driven framework, leveraging shared genetic mechanisms across complex diseases.

### Benchmarking EHR embedding methods within the EEPRS framework

To systematically evaluate the effectiveness of phenotype embeddings for genetic risk prediction, we conducted a comprehensive benchmarking analysis within the EEPRS framework, comparing five embedding methods: Word2Vec, Word2Vec_PCA, Word2Vec_ICA, GPT_PCA, and GPT_ICA. For each target trait, we first constructed a single-trait PRS using PRS-CS-auto based on external GWAS summary statistics. Next, this single-trait PRS was integrated with EHR embedding-based PRS from each embedding method using our EEPRS-Integrator, guided by the same external target trait GWAS summary statistics. The prediction accuracy of EEPRS was evaluated for 41 clinical traits available in the independent UKBB testing set (*N* = 103, 867, Supplementary Table 2). Prediction accuracy was quantified by *R*^2^ for continuous traits and the Area Under the Receiver Operating Characteristic Curve (AUC) for binary traits.

Figure 3 and Supplementary Table 3 show that EEPRS consistently improved genetic risk prediction across the 40 evaluated traits compared to the original single-trait PRS, suggesting effective integration for all five embedding methods assessed. Ovarian cancer (OvC) was excluded from the evaluation, as it was the only trait whose single-trait PRS yielded an AUC below 0.5 (AUC = 0.498 *<* 0.5).

**Figure 3:**
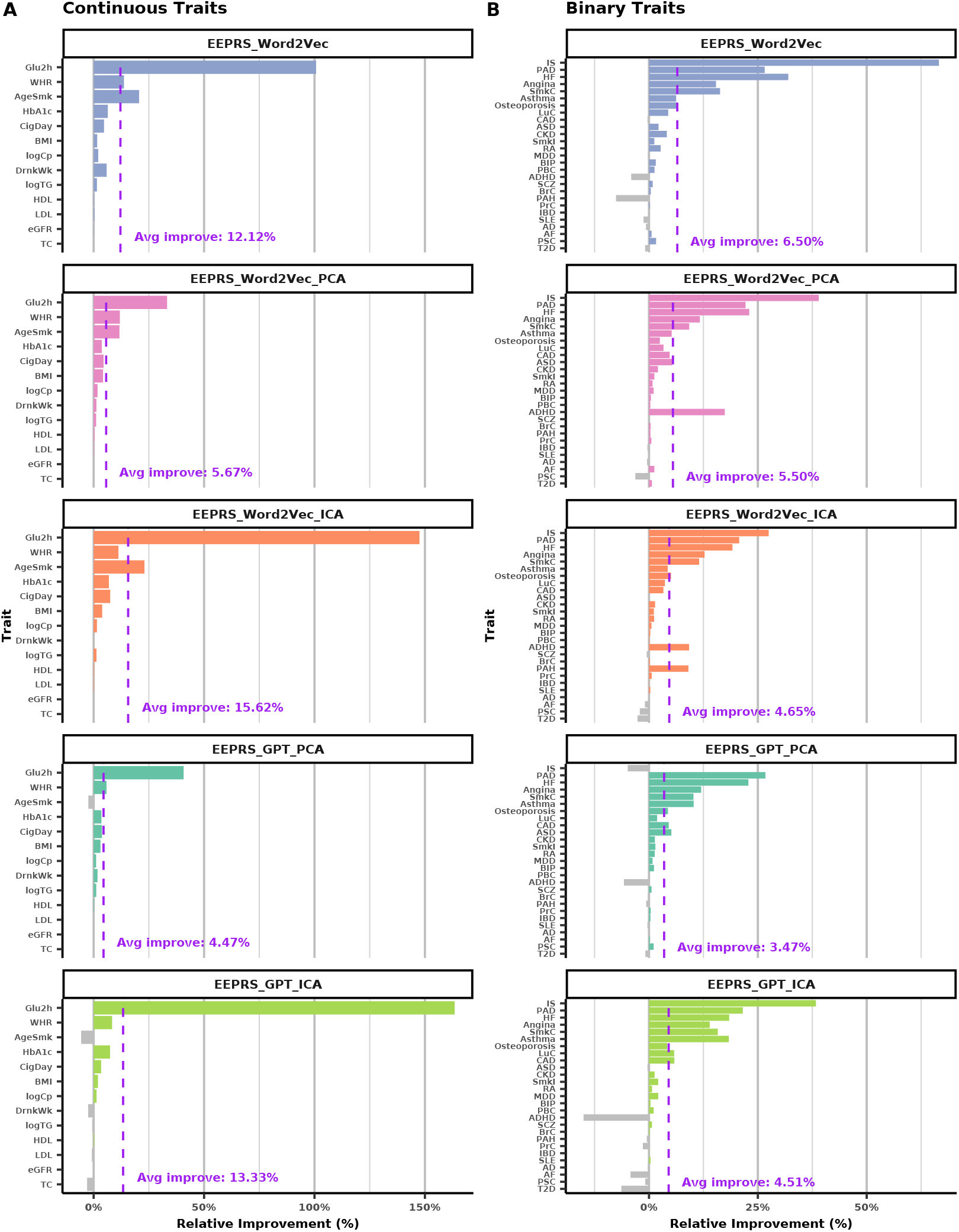
Improved prediction performance of EEPRS embedding methods versus PRSCS-auto. Bar plots illustrating relative improvements in prediction accuracy of EEPRS with five embedding methods (Word2Vec, Word2Vec_PCA, Word2Vec_ICA, GPT_PCA, and GPT_ICA) compared to PRS-CS-auto, using UKBB testing data for **A** 13 continuous traits and **B** 27 binary traits. Relative improvements are quantified as 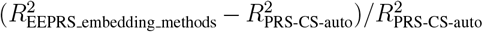 for continuous traits and 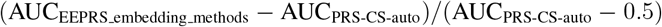 for binary traits. Horizontal colored bars represent improvements for individual traits, and grey bars indicate decreases. Vertical dashed purple lines show average improvements across traits, annotated with average improvement percentages.

Specifically, Word2Vec EEPRS achieved substantial average improvements, with an average relative increase of 12.12% in *R*^2^ for 13 continuous traits and 6.50% in AUC for 27 binary traits. Word2Vec_PCA EEPRS exhibited moderate improvements, averaging 5.67% in *R*^2^ and 5.50% in AUC. Notably, Word2Vec_ICA EEPRS provided the most improvement for continuous traits, averaging a 15.62% relative increase in *R*^2^, alongside a 4.65% improvement in AUC for binary traits. GPT_PCA EEPRS resulted in smaller relative performance gains, averaging 4.47% in *R*^2^ and 3.47% in AUC. GPT_ICA EEPRS showed significant improvements, averaging 13.33% in *R*^2^ and 4.51% in AUC. These results show variability in prediction performance improvement across embedding methods and traits, suggesting Word2Vec_ICA EEPRS effective for continuous traits and Word2Vec EEPRS effective for binary traits.

Overall, the above results demonstrate that the improved performance of EEPRS is robust across different EHR embedding methods and can be further optimized for each trait.

### Leveraging optimal EHR embeddings enhances single-trait PRS

To optimize risk prediction accuracy within EEPRS, we implemented a four-fold cross-validation procedure to identify the optimal embedding method for each trait in the UKBB testing cohort (*N* = 103, 867). Specifically, we partitioned the UKBB data into four equal subsets. For each cross-validation fold, three subsets (75%) were used as the tuning set to identify the embedding method that had the best prediction performance, quantified by *R*^2^ for continuous traits and the AUC for binary traits. The embedding method with the best performance in the tuning set was used to construct an optimized EEPRS (EEPRS_optimal), which was subsequently applied to the remaining subset (25%) designated as the testing set. This procedure was repeated across all four folds, with the selected EEPRS method presented in Supplementary Figure 6. The final prediction performance was calculated by averaging results from the four testing subsets.

As illustrated in Figure 4 and Supplementary Table 4, this optimization strategy improved prediction accuracy. EEPRS_optimal achieved notable average relative improvements of 16.05% in *R*^2^ across 13 continuous traits and 12.35% in AUC across 27 binary traits, compared to the corresponding single-trait PRS without embedding integration. These findings underscore the potential benefits of employing single-trait optimal embedding methods within the EEPRS framework, highlighting its efficacy in enhancing genetic risk prediction accuracy.

**Figure 4:**
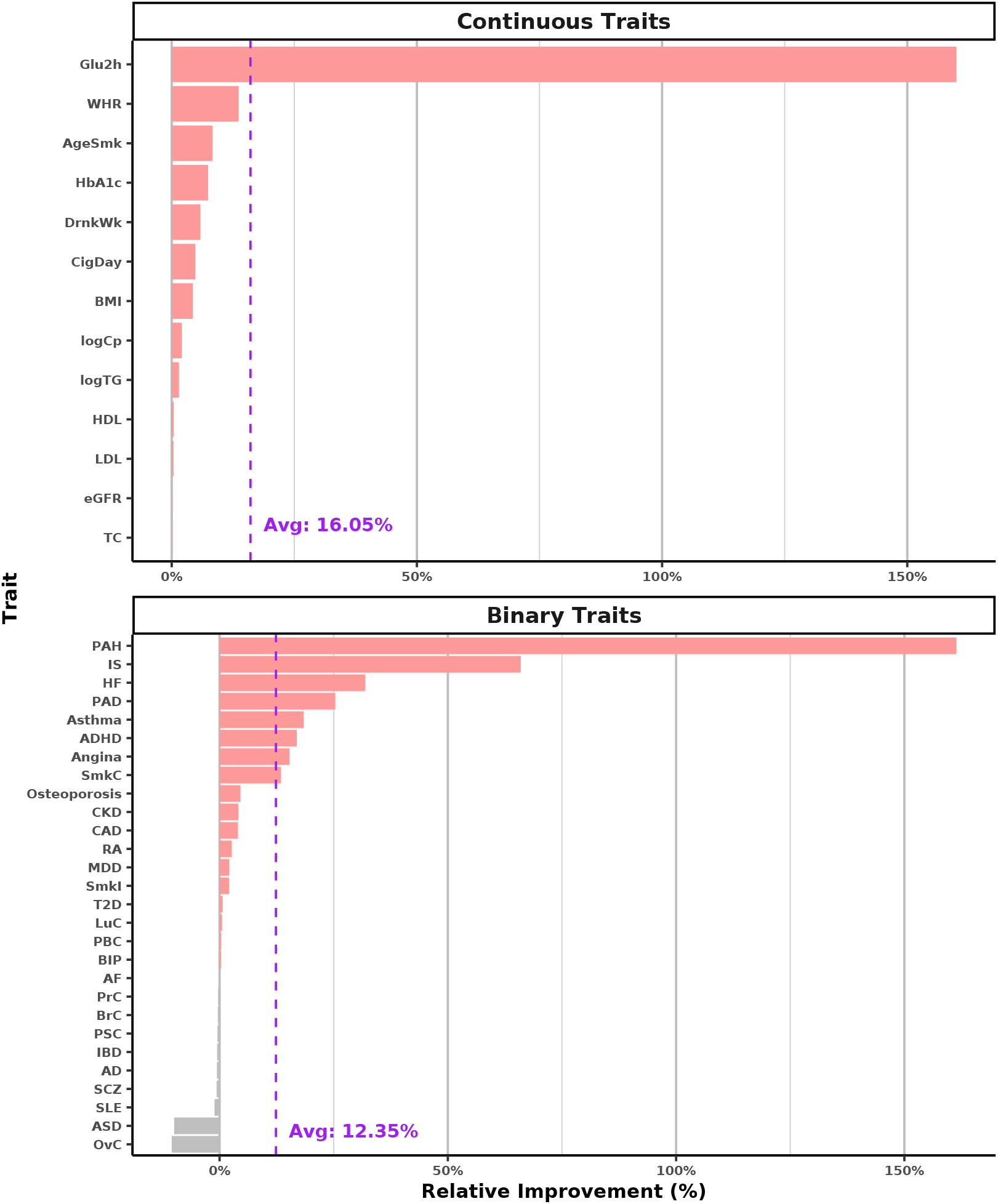
Improved prediction performance of EEPRS_optimal versus PRS-CS-auto. Bar plots illustrating relative improvements in prediction accuracy of EEPRS_optimal compared to PRS-CS-auto, evaluated using UKBB testing data via 4-fold cross-validation for 13 continuous and 27 binary traits. Relative improvements are quantified as 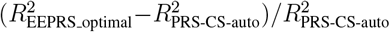 for continuous traits and 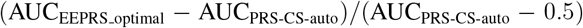 for binary traits, where *R*^2^ and AUC values represent the mean across four folds. Horizontal red bars represent improvements for individual traits, and grey bars indicate decreases. Vertical dashed purple lines indicate average improvements across traits, annotated with average improvement percentages.

Notably, pulmonary arterial hypertension (PAH), IS, HF, PAD, asthma, ADHD, and angina demonstrated the top improvements when using EEPRS_optimal compared to the single-trait PRS (PAH: 161.39%; IS: 65.96%; HF: 31.90%; PAD: 25.35%; asthma: 18.40%; ADHD: 16.91%; angina: 15.33%). These traits largely overlap with or are closely related to those comprising our identified cardiovascular cluster (logCp, T2D, PAD, IS, CAD, BMI, angina, HF), highlighting their shared clinical characteristics and genetic basis.

### Interpreting EHR embedding PRS within a PRS-based PheWAS framework

To elucidate the top improvements in the cardiovascular cluster using EEPRS and to comprehensively interpret the EHR embedding-based PRS results, we performed PRS-based PheWAS analyses to identify clinical traits associated with these PRS. We analyzed embedding methods Word2Vec_ICA and GPT_ICA, as these ICA-based embeddings included the dimensions with the highest heritability identified in previous analyses.

We then conducted a PRS-based PheWAS in the UKBB testing set (*N* = 103, 867), regressing 1,082 ICD-10 diagnostic codes on each PRS using logistic regression, adjusted for age, sex, and top 20 genetic PCs [23, 24]. After phenotype filtering, we tested associations between each PRS and 177 clinical phenotypes. We controlled for multiple testing using Benjamini–Hochberg (BH)-adjusted p-values at a significance level of 0.05 across all ICD-10 phenotypes for each PRS [65].

We categorized ICD-10 codes into 15 phenotype categories: circulatory system, dermatologic, digestive, endocrine/metabolic, genitourinary, hematopoietic, infectious diseases, mental disorders, musculoskeletal, neoplasms, neurological, pregnancy complications, respiratory, sense organs, and symptoms. To summarize PRS-based PheWAS results and identify broad patterns across clinical domains (Figure 5), we selected the smallest BH-adjusted p-value among all phenotypes within each category to represent its overall significance.

**Figure 5:**
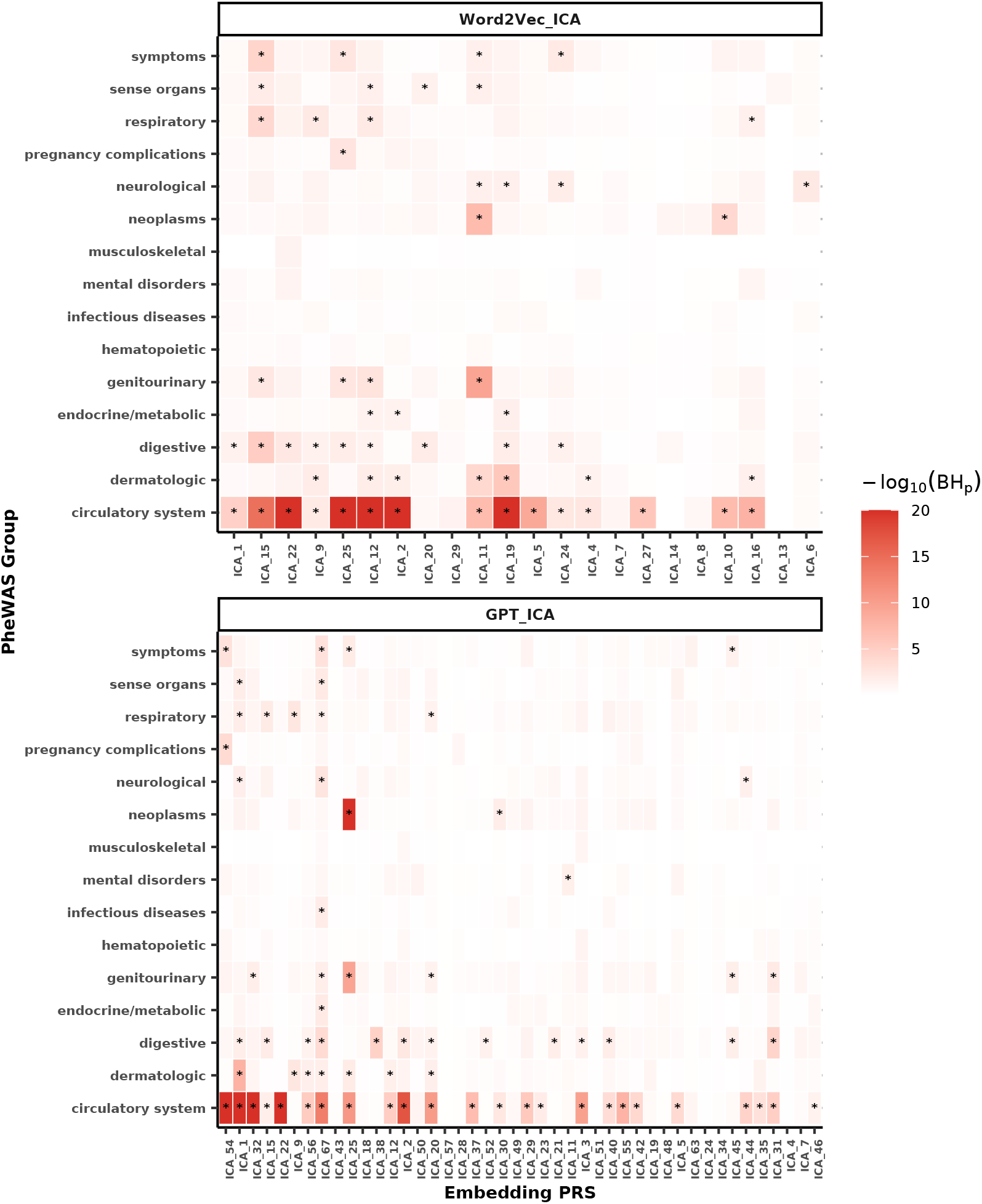
Summary of PRS-based PheWAS results. Heatmap summarizing associations between EHR embedding-based PRS from Word2Vec_ICA and GPT_ICA and disease categories identified through PRS-based PheWAS using UKBB testing data. Association strength for each category is indicated by the smallest BH-adjusted p-value across traits within that category. Color intensity reflects the strength of association, and significant associations (BH-adjusted p-values *<* 0.05) are marked by stars (*∗*).

As shown in Figure 5, distinct patterns of associations across phenotype categories indicate that that each embedding method and dimension provides unique clinical insights. Both Word2Vec ICA and GPT ICA embeddings had strong associations with circulatory system diseases (BH-adjusted p-value *<* 1 *×* 10^*−*20^). These strong associations explain why the cardiovascular cluster showed the greatest improvement in genetic risk prediction using EEPRS_optimal compared to single-trait PRS, reflecting the well-established clinical and genetic interconnectedness between cardiovascular conditions and circulatory system diseases.

GPT ICA embeddings also exhibited robust associations with neoplasms (BH-adjusted p-value *<* 1 *×* 10^*−*20^), whereas Word2Vec_ICA showed significant but slightly weaker associations in this category. Furthermore, both embedding methods consistently captured significant associations across various categories, including dermatologic, digestive, endocrine/metabolic, genitourinary, neurological, pregnancy complications, respiratory, sense organs, and symptoms. Compared to Word2Vec_AC, GPT_ICA uniquely identified significant associations with infectious diseases and mental disorders, highlighting its capability in capturing genetic susceptibility to certain clinical conditions. Conversely, neither method detected significant associations in hematopoietic or musculoskeletal categories, likely reflecting limited sample sizes for these conditions in the PheWAS analysis and indicating potential areas for methodological refinement aimed at enhancing signal detection for these specific disease categories.

Subsequently, we conducted detailed analyses of the three most heritable embedding components (Supplementary Figures 7-9). Remarkably, the top components from both methods consistently exhibited strong associations (BH-adjusted p-value *<* 1 *×* 10^*−*20^) with hypertension and essential hypertension (circulatory system diseases). Interestingly, while the top two PRS from Word2Vec_ICA showed positive associations with hypertension and essential hypertension, indicating increased genetic risk against these conditions, the top two PRS from GPT_ICA exhibited negative associations, suggesting protective genetic effects.

Additionally, both EHR embedding-based PRS identified associations with multiple clinical categories, including circulatory system diseases (atrial fibrillation and flutter, cardiac dysrhythmias), dermatologic conditions (chronic skin and leg ulcers), genitourinary conditions (urinary symptoms and disorders), neurological (CNS infections, headache syndromes), pregnancy complications (hypertension complicating pregnancy), and symptoms (syncope and collapse). Word2Vec_CA PRS uniquely captured associations with dermatologic conditions (chronic ulcers of the leg or foot), digestive diseases (ascites, dysphagia, esophageal disorders, stomach dysfunction), endocrine/metabolic diseases (fluid and electrolyte imbalance, hypovolemia, proteinuria), genitourinary conditions (pyelonephritis, urinary retention), respiratory diseases (cough, pleurisy), and sense organs (vertigo and dizziness), whereas GPT_ICA PRS showed unique associations with respiratory diseases (unspecified respiratory disorders) and sense organs (middle ear and mastoid disorders). These findings collectively suggest that different embedding methods capture diverse clinical pathways.

Overall, PRS-based PheWAS analyses demonstrated that the previously observed top improvements of EEPRS_optimal over single-trait PRS for cardiovascular clusters were driven by strong associations between EHR embedding-based PRS and circulatory system diseases. Furthermore, these analyses revealed that while different embedding methods captured shared phenotypic information, method-specific associations were also evident: GPT_ICA uniquely identified associations with infectious diseases and mental disorders, whereas Word2Vec_ICA captured a broader range of phenotypic associations. Collectively, these findings highlight the effectiveness and clinical value of our proposed PRS-based PheWAS framework for interpreting EHR embedding-based PRS.

### Incorporating EHR embeddings enhances multi-trait PRS performance

Building upon our previous findings that EEPRS_optimal can enhance single-trait PRS performance, we further investigated whether integrating EEPRS_optimal with multi-trait PRS could provide additional accuracy improvements beyond traditional multi-trait methods. Specifically, we constructed multi-trait PRS via MTAG for each pair of the 51 traits selected to represent major clinical domains using external GWAS summary statistics, with predictive evaluations remaining focused on the 41 traits available in UKBB [13, 16].

For each of the 41 target traits, we performed pairwise meta-analysis GWAS using MTAG, subsequently estimating SNP effect sizes with PRS-CS-auto for each pairwise GWAS result. This procedure generated 50 distinct multi-trait PRS per target trait, each derived from pairwise combinations with the remaining 50 traits. For each target trait, these 50 individual PRS were aggregated into a single, robust score (MTAG_PRS) using elastic net regression trained on the UKBB training dataset (*N* = 207, 734).

We then integrated EEPRS_optimal with MTAG_PRS to form an enhanced integrated model termed MTAG_EEPRS. The prediction performance of MTAG_EEPRS was evaluated using four-fold cross-validation within the UKBB testing dataset (*N* = 103, 867). In each fold, the tuning fold (75% of the testing data) identified the optimal EEPRS embedding approach, which was then combined with MTAG_PRS in the same tuning fold. The prediction performance of the resulting MTAG_EEPRS model was assessed in the corresponding testing fold (25%), and the final performance metrics were averaged across the four independent testing folds (*R*^2^ for continuous traits and AUC for binary traits).

As illustrated in Figure 6 and Supplementary Table 5, integrating EEPRS with multi-trait PRS (MTAG_EEPRS) could improve prediction accuracy, yielding notable average relative improvements of 92.48% in *R*^2^ across 13 continuous traits and 24.06% in AUC across 27 binary traits (with OvC excluded as its single-trait PRS AUC *<* 0.5), compared to the original single-trait PRS. Furthermore, when directly compared to MTAG_PRS alone (Supplementary Figure 10), MTAG_EEPRS provided incremental improvements of 1.17% in *R*^2^ for 13 continuous traits and 5.69% in AUC for 28 binary traits (with OvC included as its MTAG_PRS AUC *>* 0.5). These results suggest that EEPRS captures complementary genetic signals not fully utilized by traditional multi-trait PRS methods, underscoring the value added by integrating EEPRS into multi-trait genetic risk prediction frameworks.

**Figure 6:**
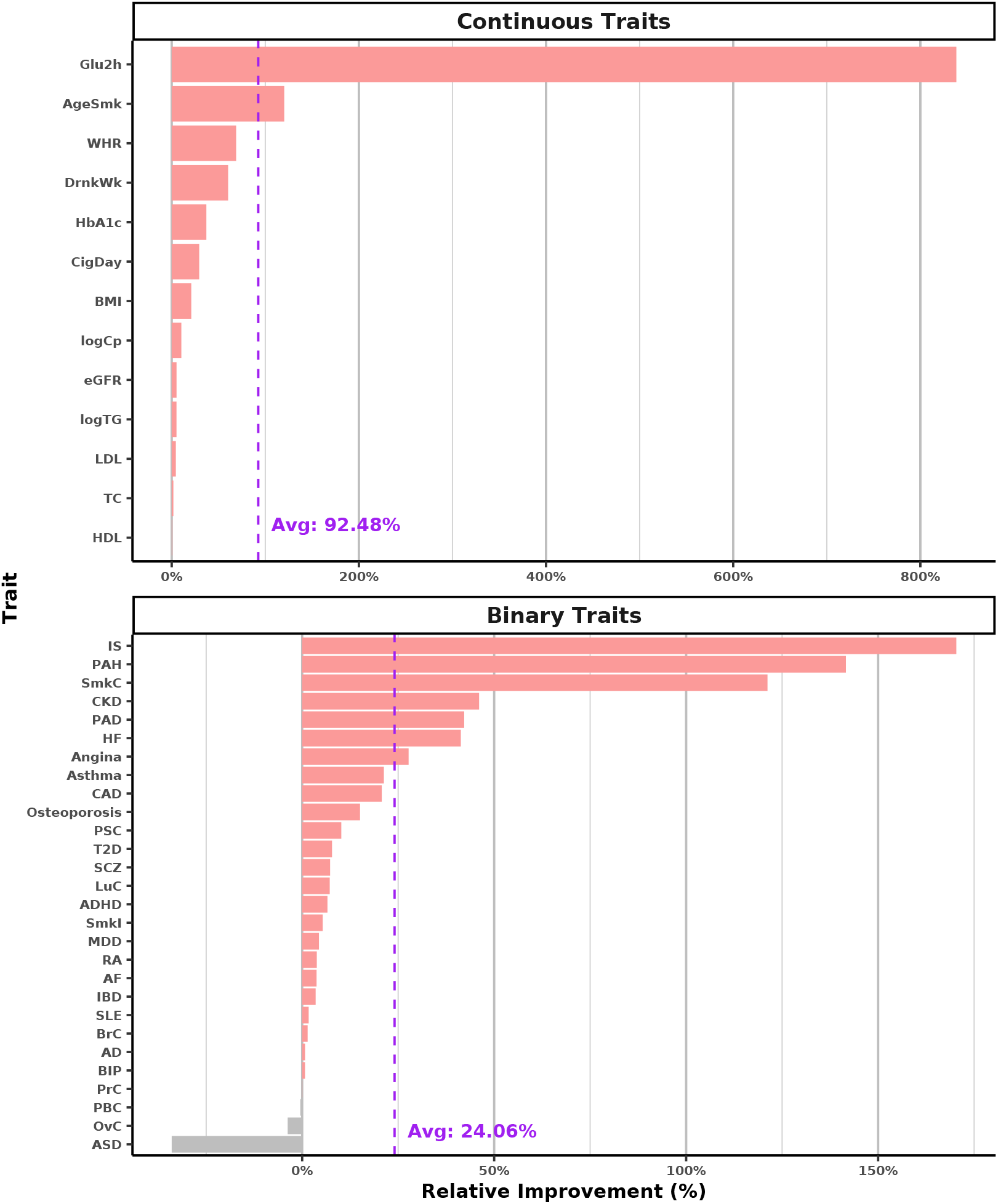
Improved prediction performance of MTAG_EEPRS versus PRS-CS-auto. Bar plots illustrating relative improvements in prediction accuracy of MTAG_EEPRS compared to PRS-CS-auto, evaluated using UKBB testing data via 4-fold cross-validation for 13 continuous and 27 binary traits. Relative improvements are quantified as 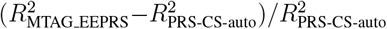 for continuous traits and 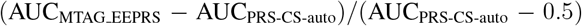 for binary traits, where *R*^2^ and AUC values represent the mean across four folds. Horizontal red bars represent improvements for individual traits, and grey bars indicate decreases. Vertical dashed purple lines indicate average improvements across traits, annotated with average improvement percentages.

## Discussion

In this study, we introduced EEPRS, a novel PRS framework developed to systematically integrate EHR embeddings with single-trait and multi-trait PRS to enhance genetic risk prediction. EEPRS leverages the existing embedding methods, including Word2Vec and GPT, to effectively convert complex, high-dimensional EHRs into informative, lower-dimensional phenotypic vector representations. GWAS performed on these embeddings demonstrated statistically significant heritability and strong genetic correlations with diverse traits. Hierarchical clustering based on genetic correlation profiles further identified a distinct cardiovascular cluster.

We evaluated EEPRS on 41 clinical traits from the UK Biobank and showed that it consistently outperformed traditional single-trait PRS methods, particularly within the cardiovascular cluster. Validation through PRS-based phenome-wide association studies (PRS-PheWAS) further confirmed robust associations between EHR embedding-based PRS and circulatory system diseases.

We further improved prediction accuracy by employing a data-adaptive cross-validation approach to identify the optimal embedding method for each trait, additionally integrating multitrait PRS. Although our current analysis utilized individual-level tuning data for optimal embedding selection and multi-trait integration, EEPRS-Integrator is capable of executing the complete pipeline solely based on GWAS summary statistics. This flexibility underscores EEPRS’s potential as a widely accessible tool for improving genetic risk prediction through integration of EHR embeddings.

We acknowledge that the performance of EEPRS depends on the chosen embedding methods. While our findings indicate consistent performance improvements with Word2Vec and GPT-based embeddings, future research is needed to explore other embedding approaches, such as FastText [66], BERT [67], and other LLMs [68], to potentially uncover additional genetic insights from diverse EHR representations.

Moreover, utilizing other PRS methods could also improve EEPRS performance. Single-trait PRS methods such as SDPR [8] and SBayesRC [9] may further enhance prediction accuracy and robustness. Additionally, multi-trait analyses [12, 13, 14, 15] and multi-population approaches [69, 70] may improve the generalizability and clinical utility of PRS predictions.

Moving forward, given the flexibility of EEPRS, it can be extended beyond leveraging EHR embeddings to include other data modalities such as spirograms, photoplethysmograms, and electrocardiograms [18]. This could help provide a more comprehensive genetic risk assessment integrating diverse data modalities.

## Methods

### EEPRS framework

#### Step 1: Generate EHR-based phenotype embeddings

This section describes the procedure for constructing phenotype embeddings from EHRs using ICD-10 codes, specifically addressing: (i) data preprocessing, (ii) ICD-10 code embedding, (iii) individual embedding, and (iv) dimension reduction.

##### (i) Data preprocessing

We extract each individual’s longitudinal ICD-10 code history from the UKBB [1], ordered by hospital admission dates. Each individual’s sequence of clinical descriptions associated with diagnosis codes is treated as a “sentence,” with each ICD-10 code acting as a token representing a distinct medical condition.

##### (ii) ICD-10 code embedding

We construct dense vector representations for each ICD-10 code using two complementary approaches:

- **Word2Vec-based descriptions:** Following the PERADIGM framework [71], we compile all ICD-10 code descriptions and train a Word2Vec model (CBOW architecture) to learn word embeddings [20]. For each ICD-10 code, we tokenize its clinical description, remove stopwords and punctuation, and average the corresponding word embeddings to produce the code-level representation.
- **GPT-based descriptions:** We prompt GPT-o1 to generate concise, plain-language descriptions for each ICD-10 code [21]. Each word in the generated description is embedded using the text-embedding-3-large, and the code embedding is computed as the average of these word embeddings, capturing semantic context from the language model.

##### (iii) Individual embedding

Each individual *i* is represented by the average of all embedded ICD-10 codes in his/her record:

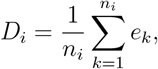

where *n*_*i*_ is the number of ICD-10 codes for individual *i*, and *e*_*k*_ denotes the embedding of the *k*-th code. This unweighted average yields a compact summary of an individual’s health profile.

##### (iv) Dimension reduction

To further compress the embeddings, we apply dimensionality reduction techniques including PCA [26] and ICA [27]. Dimensionality reduction mitigates redundancy and noise inherent in high-dimensional embeddings, thus isolating the most informative and independent features for downstream analyses. We retain principal components explaining at least 80% of the total variance, yielding reduced representations: Word2Vec_PCA, Word2Vec_ICA, GPT_PCA, and GPT_ICA.

#### Step 2: Perform EHR embedding-based GWAS

Before performing GWAS, we apply quantile normalization separately to each embedding dimension derived from all embedding methods (Word2Vec, Word2Vec_PCA, Word2Vec_ICA, GPT, GPT_PCA, GPT_ICA). This step ensures that each dimension approximately follows a normal distribution, satisfying the assumptions required for GWAS.

Each normalized embedding dimension is treated as an individual phenotype in GWAS. We then perform marginal linear regression analyses across all HapMap3 SNPs (*S* = 1, 297, 431). The regression model includes age, sex, and top 20 genetic PCs as covariates to account for population stratification and other potential confounders:

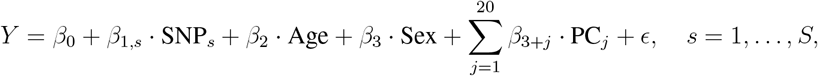

where *Y* denotes a normalized embedding dimension, *β* values are the regression coefficients, and *ϵ* represents the residual error. GWAS analyses are performed using PLINK2 [72].

#### Step 3: Derive EHR embedding-based PRS

Using EHR embedding-based GWAS summary statistics, we compute PRS with PRS-CS-auto [6], a Bayesian method that estimates SNP effect sizes by adaptively optimizing the shrinkage parameter without manual tuning. Throughout this process, linkage disequilibrium (LD) reference panels constructed from the 1000 Genomes Project Phase 3 samples [73] are employed. The PRS for individual *i* is then calculated as:

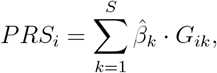

where 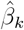 is the posterior effect size estimated by PRS-CS-auto for SNP *k*, based on the EHR embedding-based GWAS; *G*_*ik*_ denotes the genotype dosage of SNP *k* for individual *i*; and *S* is the total number of SNPs included in the analysis. The resulting PRS captures individual-level genetic predispositions based on the multidimensional phenotypic structure learned from EHR embeddings.

#### Step 4: Interpret EHR embedding-based PRS through PRS-based PheWAS

We interpret the EHR embedding-based PRS using a PRS-based PheWAS framework, assessing associations between each EHR embedding-based PRS (predictor) and ICD-10 code-derived phenotypes (outcomes) using the PheWAS R package [24]. For each PRS–phenotype pair, we perform logistic regression adjusted for age, sex, and the top 20 genetic PCs to account for population stratification and potential confounders.

The regression model for phenotype *Y* and PRS predictor is defined as:

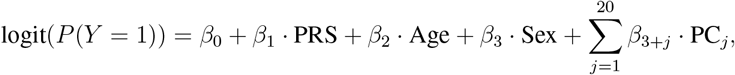

where logit(*P* (*Y* = 1)) denotes the log-odds of having the phenotype, *β*_1_ quantifies the effect of the PRS, and the remaining coefficients adjust for covariates.

To correct for multiple testing, we apply the BH procedure to control the FDR [65]:

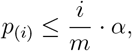

where *p*_(*i*)_ is the *i*-th smallest p-value, *m* is the total number of tests, and *α* is the desired FDR level (*α* = 0.05).

#### Step 5: Integrating EHR embedding-based PRS into the EEPRS framework

This section outlines the procedure for integrating EHR embedding-based PRS with single-trait PRS within the EEPRS framework and describes its extension to incorporate multi-trait PRS.

##### (i) EEPRS-Integrator

Building on our previously developed PRS integration framework, MIXPRS [25], we developed EEPRS-Integrator to construct EEPRS by integrating EHR embeddingbased PRS with single-trait PRS. EEPRS-Integrator requires only GWAS summary statistics, leveraging data fission principles, and consists of four steps:

###### Step 1: EHR embedding selection

We identify embedding GWAS results that are genetically correlated with the target trait, as assessed using LDSC [64]. Only embeddings showing statistically significant genetic correlation are selected for subsequent integration (*p*-value *<* 0.05).

###### Step 2: Target trait GWAS subsampling

GWAS summary statistics for the target trait are split into statistically independent training and tuning GWAS using the GWAS subsampling procedure from MIXPRS. LD pruning (pairwise *r*^2^ *<* 0.5 within 250 kb windows) is applied to mitigate LD mismatch and reduce computational burden in downstream analysis [74].

###### Step 3: Estimating PRS combination weights

This step consists of two sub-steps:

- **Step 3.1: PRS coefficient estimation:** PRS-CS-auto is applied independently to the subsampled training GWAS summary statistics for the target trait and each selected embedding GWAS (after additional LD pruning), generating LD-pruned PRS beta coefficients.
- **Step 3.2: Combination weight determination:** In contrast to the non-negative least squares approach used in MIXPRS, we employ linear regression to estimate optimal combination weights. This allows for negative weights, accommodating embeddings that may be negatively associated with the target trait. Only embeddings selected in Step1 are included to ensure robustness. Final weights are estimated using the subsampled tuning GWAS summary statistics and the calculated LD-pruned PRS beta coefficients.

###### Step 4: Derivation of EEPRS

Complete GWAS summary statistics for the target trait and selected embeddings are re-analyzed using PRS-CS-auto to generate final PRS beta coefficients. These beta coefficients are then integrated using the weights from Step 3 to derive the final EEPRS. This integrated score captures complementary genetic signals, enhancing predictive accuracy across a range of complex traits.

##### (ii) Incorporating MTAG_PRS

We further extend the EEPRS framework by incorporating multi-trait genetic information using MTAG [13, 16]. For each trait pair, MTAG is applied to produce two single-trait GWAS outputs, each capturing shared genetic signals while preserving trait specificity. For each pair, we retain the MTAG GWAS results corresponding to the target trait and estimate SNP effect sizes using PRS-CS-auto.

The resulting PRS scores are combined using elastic net regression on individual-level training data to produce an aggregate multi-trait PRS (MTAG_PRS). Finally, EEPRS and MTAG_PRS are linearly combined using individual-level tuning data to generate the final integrated score (MTAG_EEPRS). This extension further improves prediction by leveraging genetic correlations across related traits.

### Real-data analysis

We confirm that this research complies with all relevant ethical regulations. Participants from the UK Biobank provided written informed consent (further details available at https://www.ukbiobank.ac.uk/learn-more-about-uk-biobank/governance).

### GWAS summary statistics

We compiled GWAS summary statistics for 51 traits (23 continuous traits and 28 binary traits) from European-ancestry populations across multiple consortia [29, 30, 31, 32, 33, 34, 35, 36, 37, 38, 39, 40, 41, 42, 43, 44, 45, 46, 47, 48, 49, 50, 51, 52, 53, 54, 55, 56, 57, 58, 59, 60, 61, 62, 63]. All selected GWAS explicitly excluded UKBB participants to avoid sample overlap. Detailed information is provided in Supplementary Table 1.

### UKBB data

We classified UKBB individuals into five super-populations by performing PCA jointly with the 1000 Genomes Project reference samples and subsequently training a random forest classifier based on the top ten PCs to assign ancestry labels. Population counts were: 311,600 for Europeans, 2,091 for East Asians, 6,829 for Africans, 7,857 for South Asians, and 635 for Admixed Americans. This study focuses exclusively on the European population.

Among the 51 traits with available GWAS summary statistics, 41 were also included in the UKBB cohort. We obtained data on these 41 traits (13 continuous traits and 28 binary traits) from UKBB participants; detailed descriptions are provided in Supplementary Table 2.

### Evaluation metrics

For continuous traits, predictive accuracy was assessed using *R*^2^:

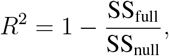

where SS_null_ and SS_full_ denote residual sums of squares from the null model (including age, sex, and top 20 genetic PCs) and the full model (additionally including PRS), respectively. To compare between methods, we defined the relative improvement of method B over method A as:

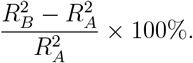

For binary traits, predictive accuracy was assessed using logistic regression, quantified by the area under the receiver operating characteristic curve (AUC). The relative improvement of method B over method A was calculated as:

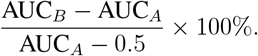

## Data availability

We will publicly release our data upon publication, including Embedding GWAS summary statistics derived from five embedding approaches (Word2Vec, Word2Vec_PCA, Word2Vec_ICA, GPT_PCA, GPT_ICA), generated using data from the UK Biobank European training population (*N* = 207, 734).

## Code availability

The code for all real-data analyses presented in this paper is publicly available on GitHub at https://github.com/LeqiXu/EEPRS_analysis. The pipeline implementing the proposed EEPRS framework is available at https://github.com/YCSGP/EEPRS.

Implementations of other methods are accessible via their respective repositories: PRS-CS-auto at https://github.com/getian107/PRScs, and the PheWAS R package at https://github.com/PheWAS/PheWAS.

## Acknowledgments

This research was supported in part by NIH grant R01 HG012735 and the OpenAI Research Access Program. Analyses were performed using data from the UK Biobank Resource (application number 29900). We thank all UK Biobank participants, whose invaluable contributions made this study possible. We would like to thank Dr. Andrew T. DeWan for his support and for facilitating access to the dbGaP dataset (accession number 32518) through an approved application on behalf of his laboratory. The data were used in this study under authorized access by a designated lab member (J.H.).

## Author contributions

H.Z. and L.X. conceived and designed the study. L.X. developed the statistical methodology and algorithm, implemented the EEPRS pipeline, and performed all primary data analyses under the supervision of H.Z. W.Z. and T.L. generated EHR embedding data. J.H. assisted with data analysis and interpretation. Y.L., J.Z., and G.W. contributed to methodological discussions and the interpretation of findings. L.X. drafted the manuscript. H.Z., W.Z., J.H., Y.L., J.Z., G.W., and T.L. provided critical revisions. All authors reviewed, edited, and approved the final manuscript.

