## Supplemental Information for "Improving polygenic risk prediction performance through integrating electronic health records by phenotype embedding"

### 1 Supplemental Figures

**Supplementary Figure 1. Overview of the EEPRS-Integrator method.** EEPRS-Integrator integrates EHR embeddings into PRS through a structured four-step workflow: (1) selection of EHR embeddings based on genetic correlation; (2) subsampling of target trait GWAS summary statistics by combining LD-pruned SNPs with identity covariance matrices, generating subsampled training and tuning datasets; (3) estimation of PRS combining weights using LD-pruned, selected EHR embeddings for PRS construction and repeated subsampled GWAS datasets, followed by linear regression to determine optimal weights; and (4) final integration step, where PRS is recalculated using full SNPs on selected EHR embeddings and target trait GWAS statistics, combined with previously derived optimal weights to produce the EEPRS. This figure was created with BioRender.

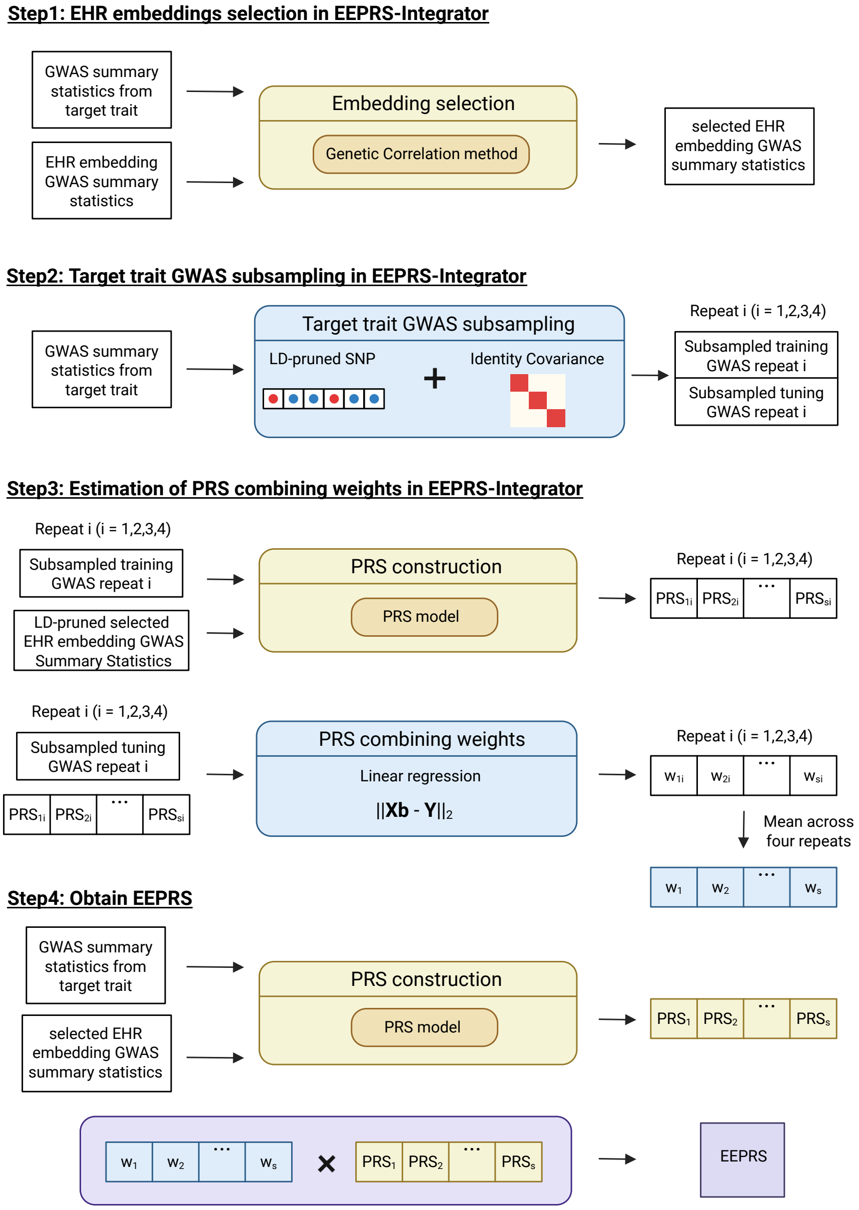

**Supplementary Figure 2. GWAS Heritability of Embedding Dimensions Across Methods.** Violin plots illustrating the distribution of GWAS heritability estimates calculated from HapMap3 SNPs for embedding dimensions derived using five embedding methods (Word2Vec, Word2Vec_PCA, Word2Vec_ICA, GPT_PCA, and GPT_ICA). Each dot represents an individual embedding dimension, with filled dots indicating dimensions that achieved statistical significance (TRUE) and open dots representing non-significant dimensions (FALSE). Horizontal lines denote median heritability values for each embedding approach.

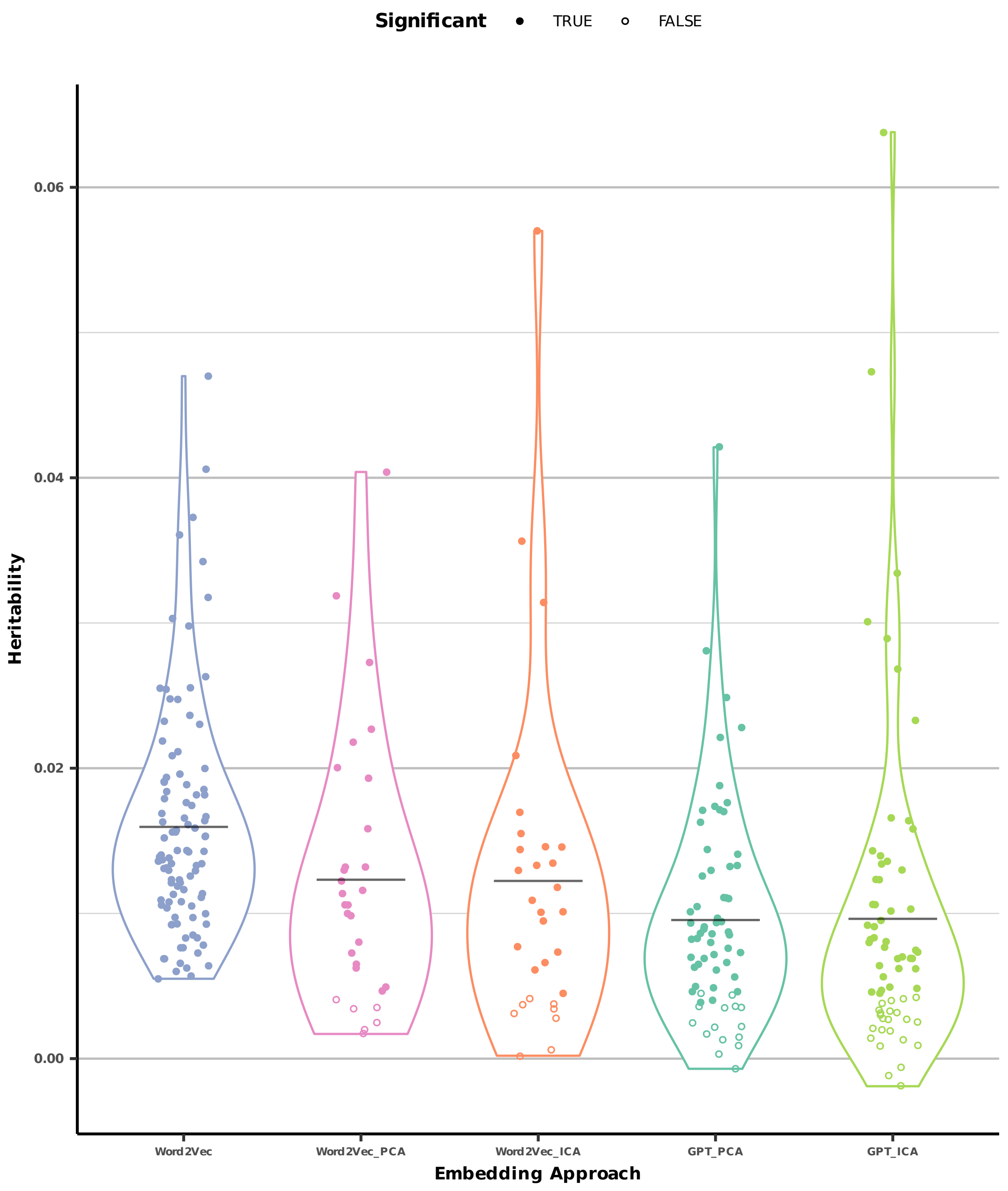

**Supplementary Figure 3. Genetic correlations between heritable Word2Vec_PCA and Word2Vec_ICA embeddings and 51 complex traits.** Heatmap illustrating genetic correlations between GWAS of heritable Word2Vec_PCA and Word2Vec_ICA embeddings and 51 complex traits. Positive correlations are indicated in shades of red, and negative correlations in shades of blue, with color intensity reflecting correlation strength. Significant correlations (BH-adjusted p-value $< 0.05$) are marked by stars ($*$).

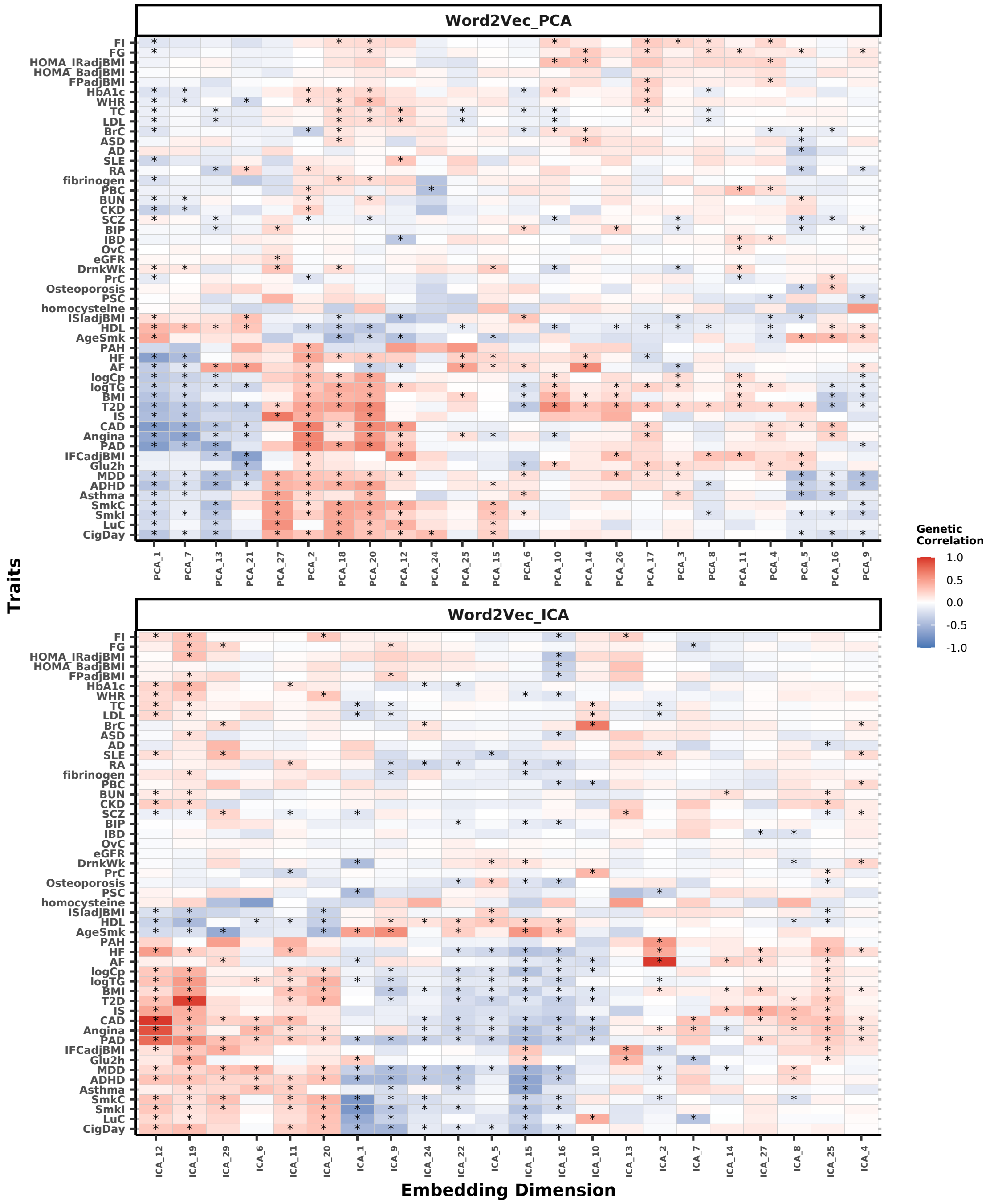

**Supplementary Figure 4. Genetic correlations between heritable GPT_PCA and GPT_ICA embeddings and 51 complex traits.** Heatmap illustrating genetic correlations between GWAS of heritable GPT_PCA and GPT_ICA embeddings and 51 complex traits. Positive correlations are indicated in shades of red, and negative correlations in shades of blue, with color intensity reflecting correlation strength. Significant correlations (BH-adjusted p-value $< 0.05$) are marked by stars ($*$).

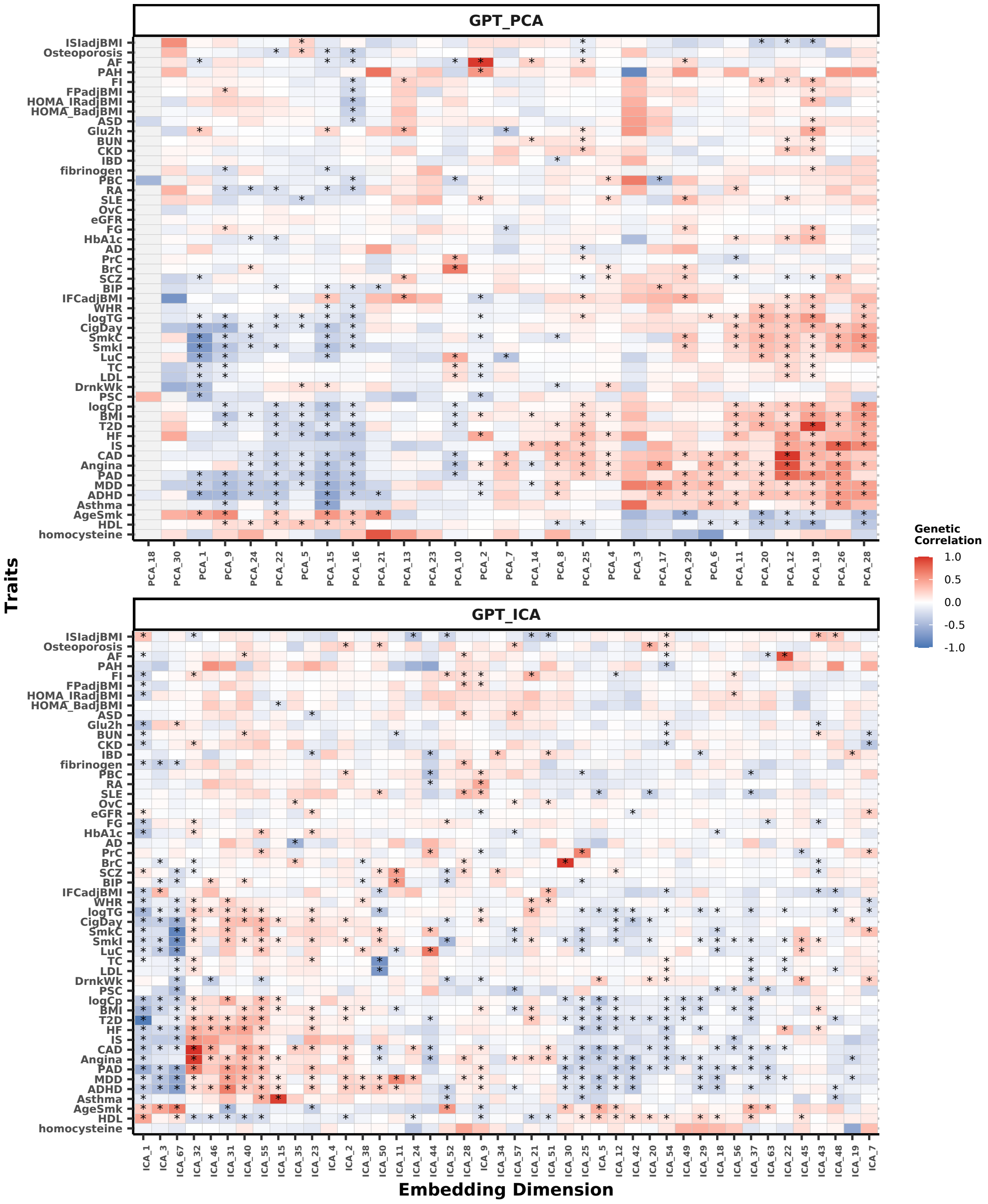

**Supplementary Figure 5. Clustering of 51 complex traits based on their genetic correlation profiles with five embedding methods.** Heatmap showing cluster assignments of 51 complex traits according to their genetic correlations with embeddings from five methods (GPT_ICA, GPT_PCA, Word2Vec, Word2Vec_ICA, and Word2Vec_PCA). Traits are grouped using unsupervised clustering based on their correlation profiles, and colors indicate cluster membership.

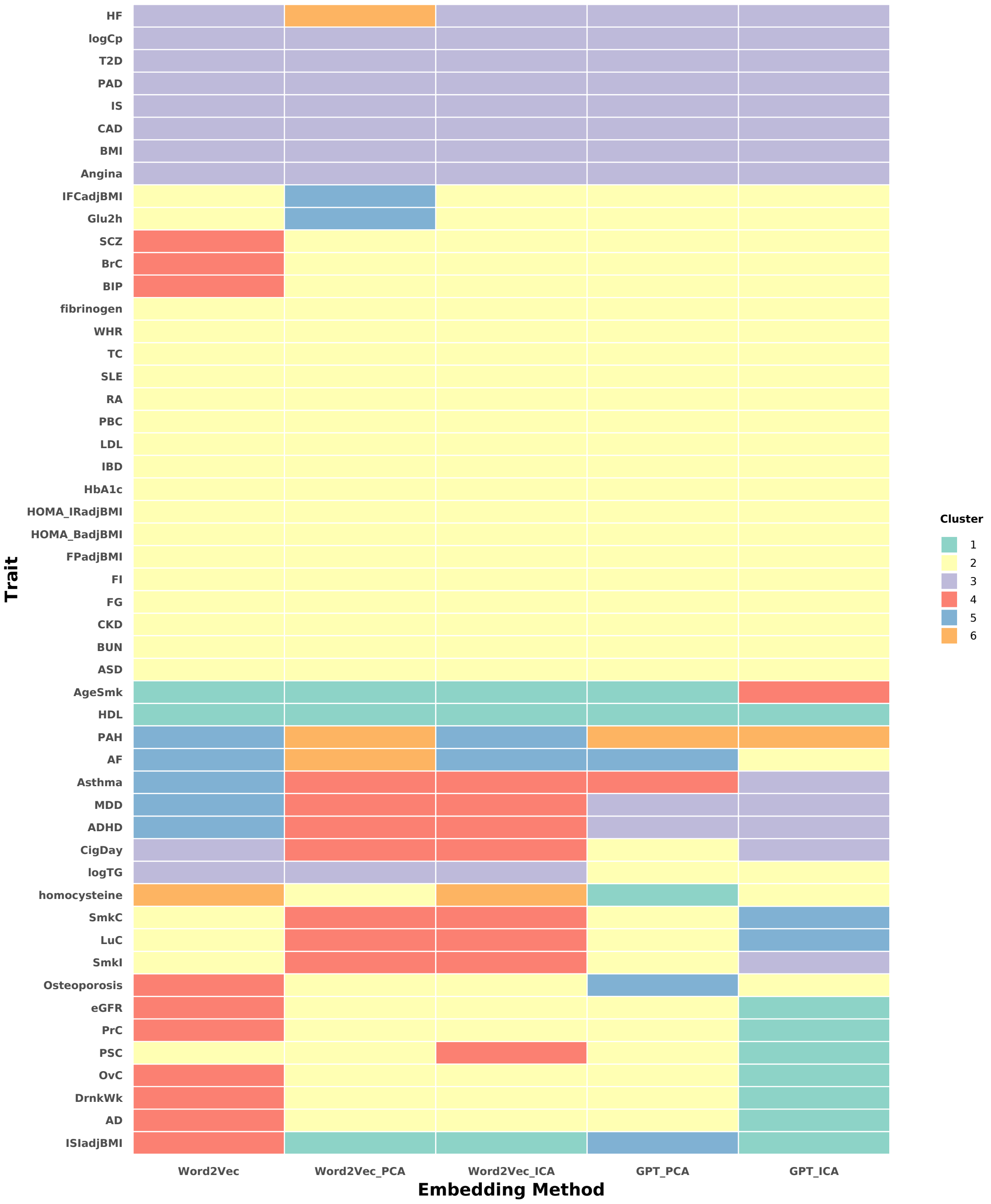

**Supplementary Figure 6. EEPRS optimal method selection across traits and folds.** The stacked bar plot illustrates the optimal EEPRS method selected across 41 traits evaluated over four cross-validation folds. Compared EEPRS methods include Word2Vec, Word2Vec_PCA, Word2Vec_ICA, GPT_PCA, GPT_ICA. Each bar segment shows the selected optimal EEPRS method for each fold.

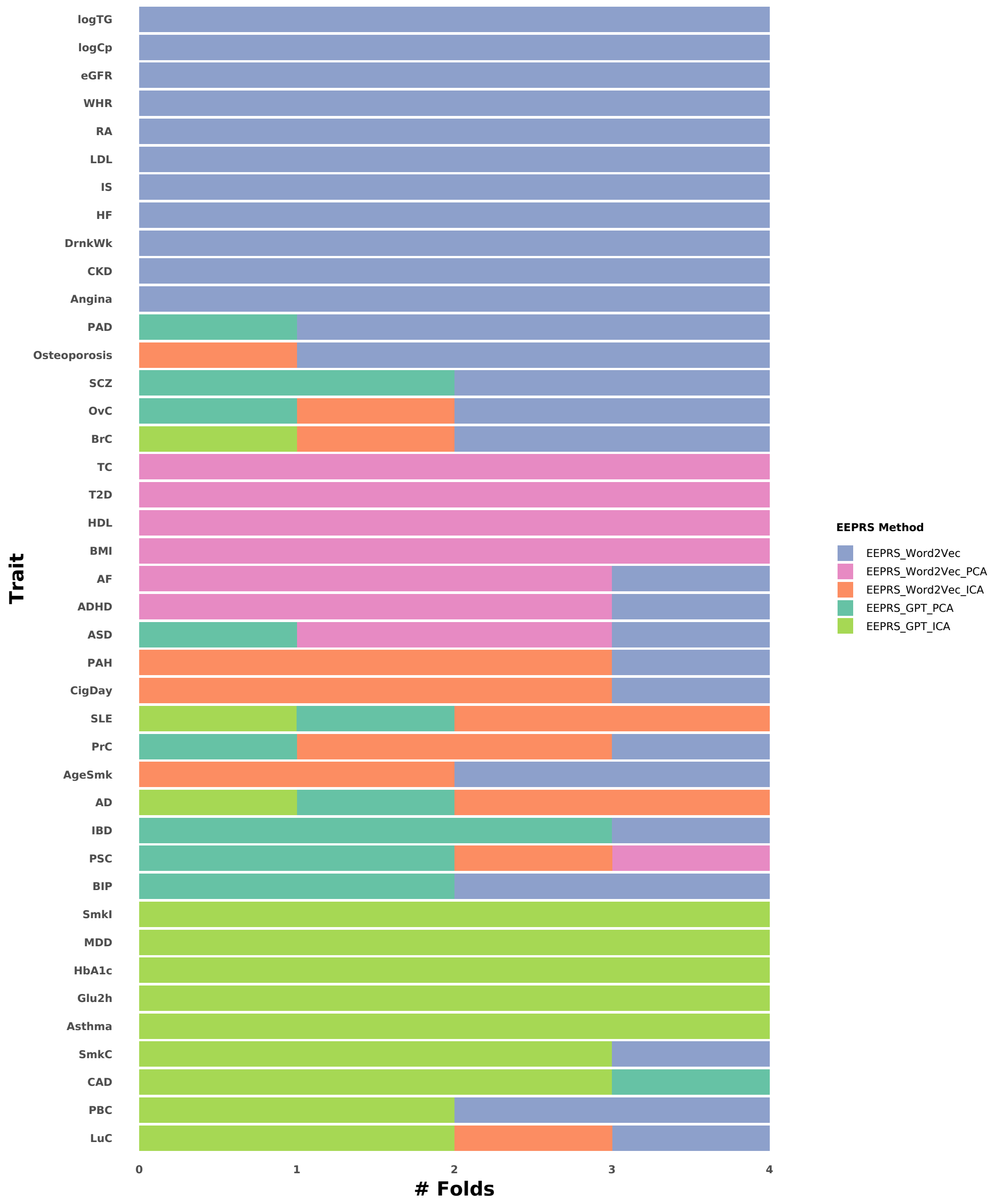

**Supplementary Figure 7. PRS-PheWAS results for the most heritable embeddings.** Scatter plot illustrating associations between PRS derived from the most heritable embeddings from Word2Vec_ICA and GPT_ICA and traits across multiple disease categories identified through PRS-based PheWAS using UKBB testing data. Triangle orientation indicates the direction of association (upward triangles for positive associations; downward triangles for negative associations). The horizontal purple line marks the significance threshold (BH-adjusted p-value $< 0.05$). Traits positioned to the right of this line exhibit significant associations.

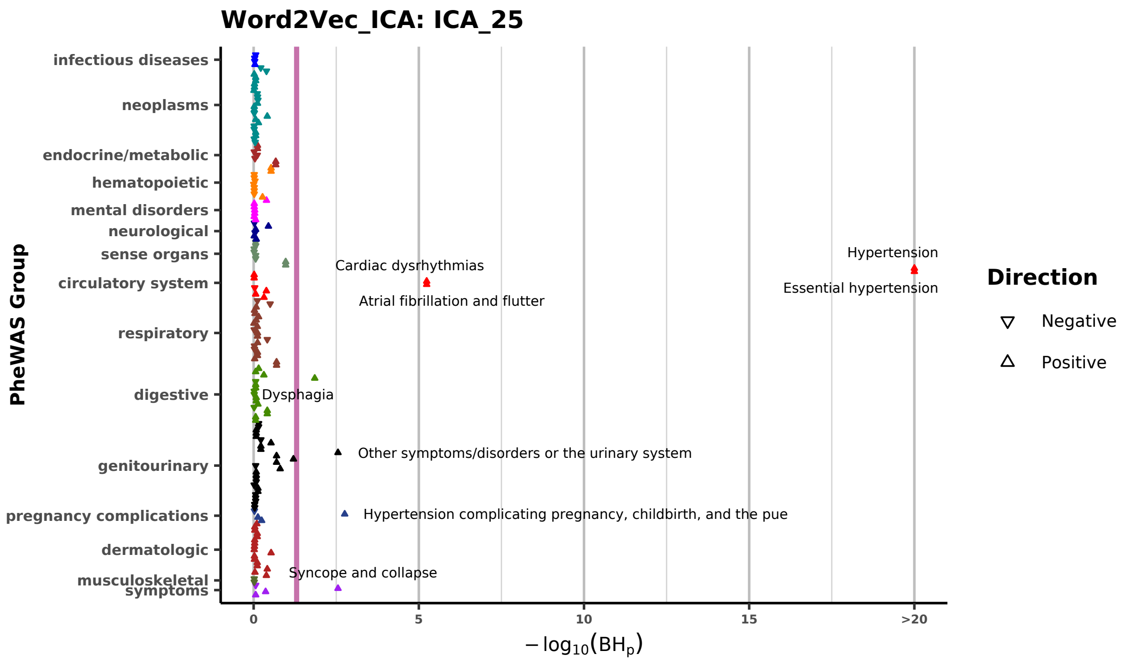

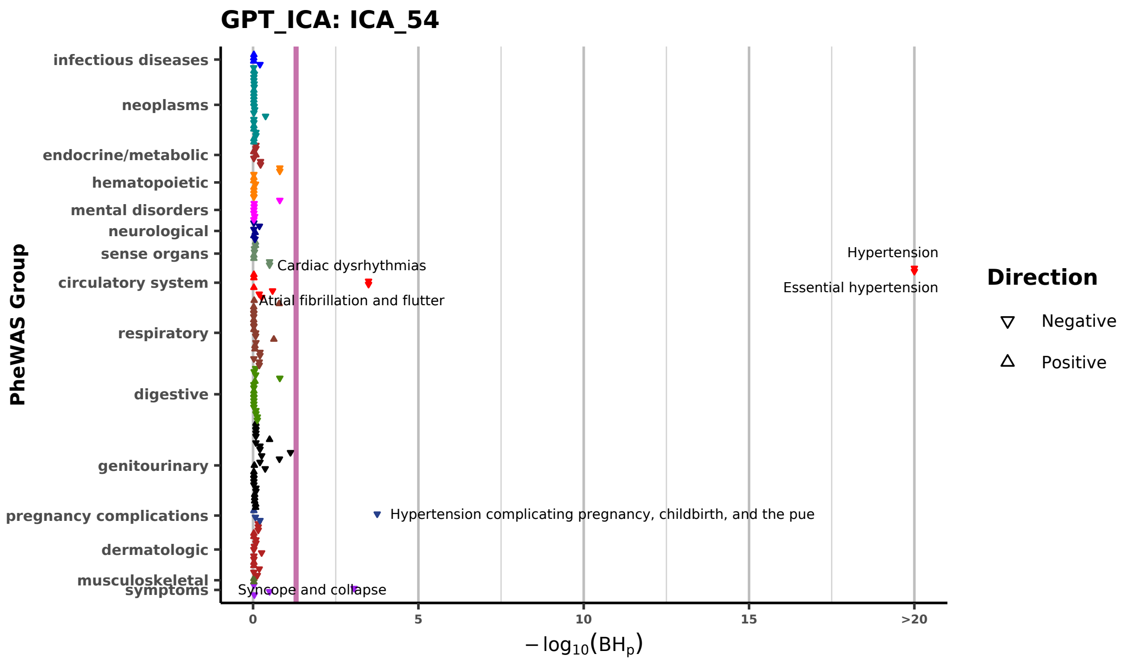

**Supplementary Figure 8. PRS-PheWAS results for embeddings with the second-highest heritability.** Scatter plot illustrating associations between PRS derived from embeddings with the second-highest heritability obtained via Word2Vec_ICA and GPT_ICA and traits across multiple disease categories identified through PRS-based PheWAS using UKBB testing data. Triangle orientation indicates the direction of association (upward triangles for positive associations; downward triangles for negative associations). The horizontal purple line marks the significance threshold (BH-adjusted $p$-value $< 0.05$). Traits positioned to the right of this line exhibit significant associations.

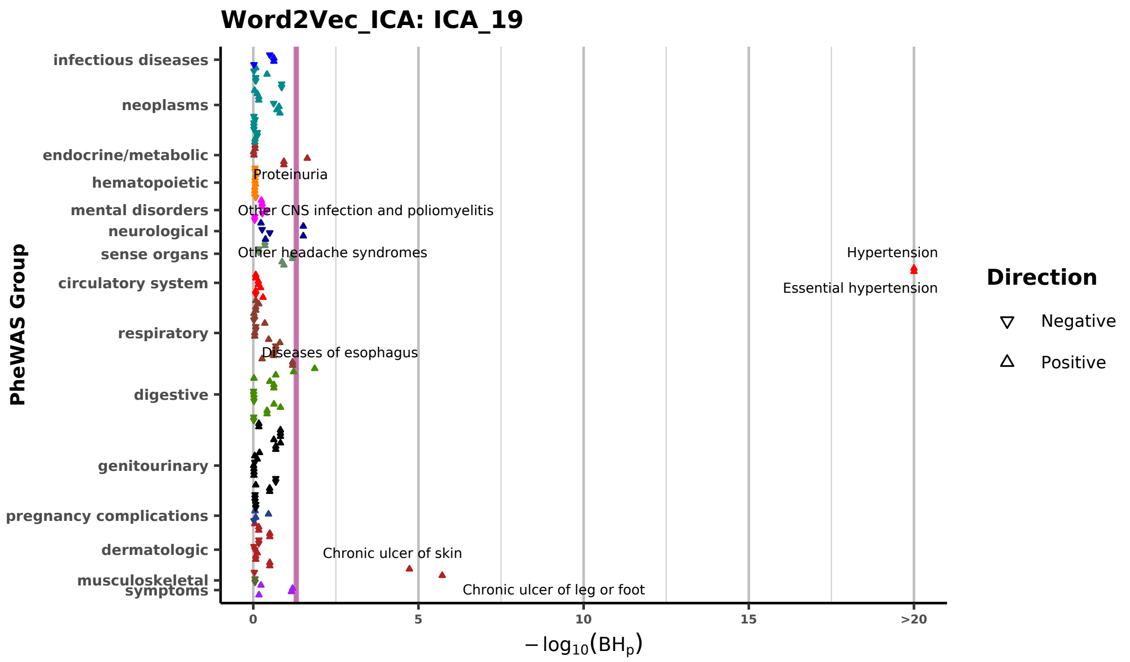

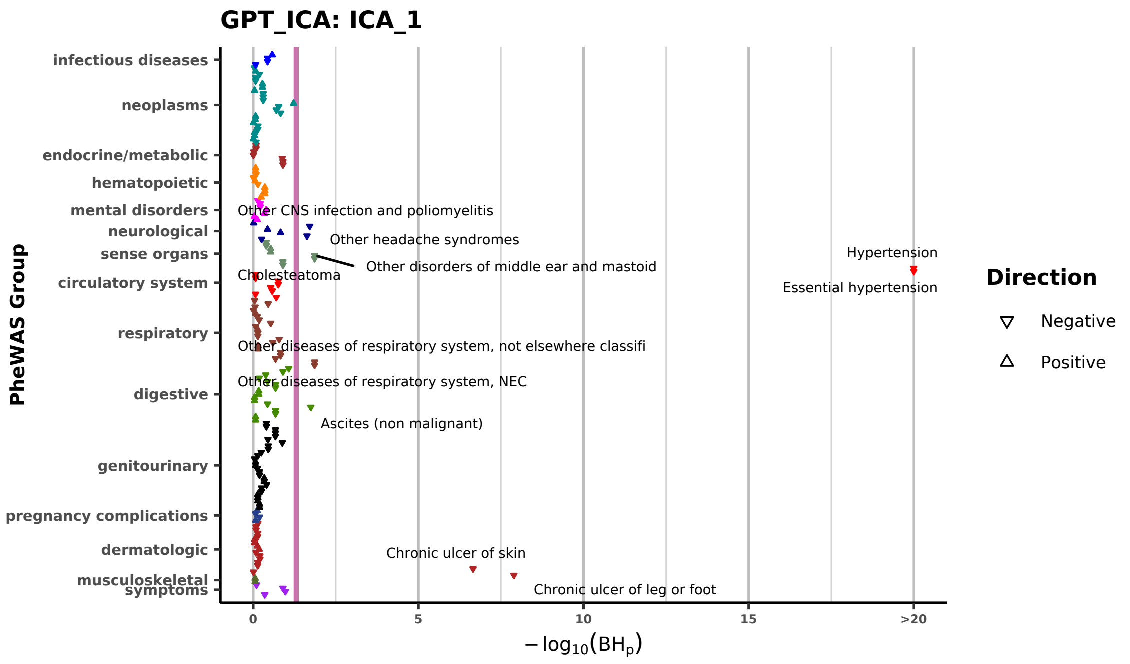

**Supplementary Figure 9. PRS-PheWAS results for embeddings with the third-highest heritability.** Scatter plot illustrating associations between PRS derived from embeddings with the third-highest heritability obtained via Word2Vec_ICA and GPT_ICA and traits across multiple disease categories identified through PRS-based PheWAS using UKBB testing data. Triangle orientation indicates the direction of association (upward triangles for positive associations; downward triangles for negative associations). The horizontal purple line marks the significance threshold (BH-adjusted $p$-value $< 0.05$). Traits positioned to the right of this line exhibit significant associations.

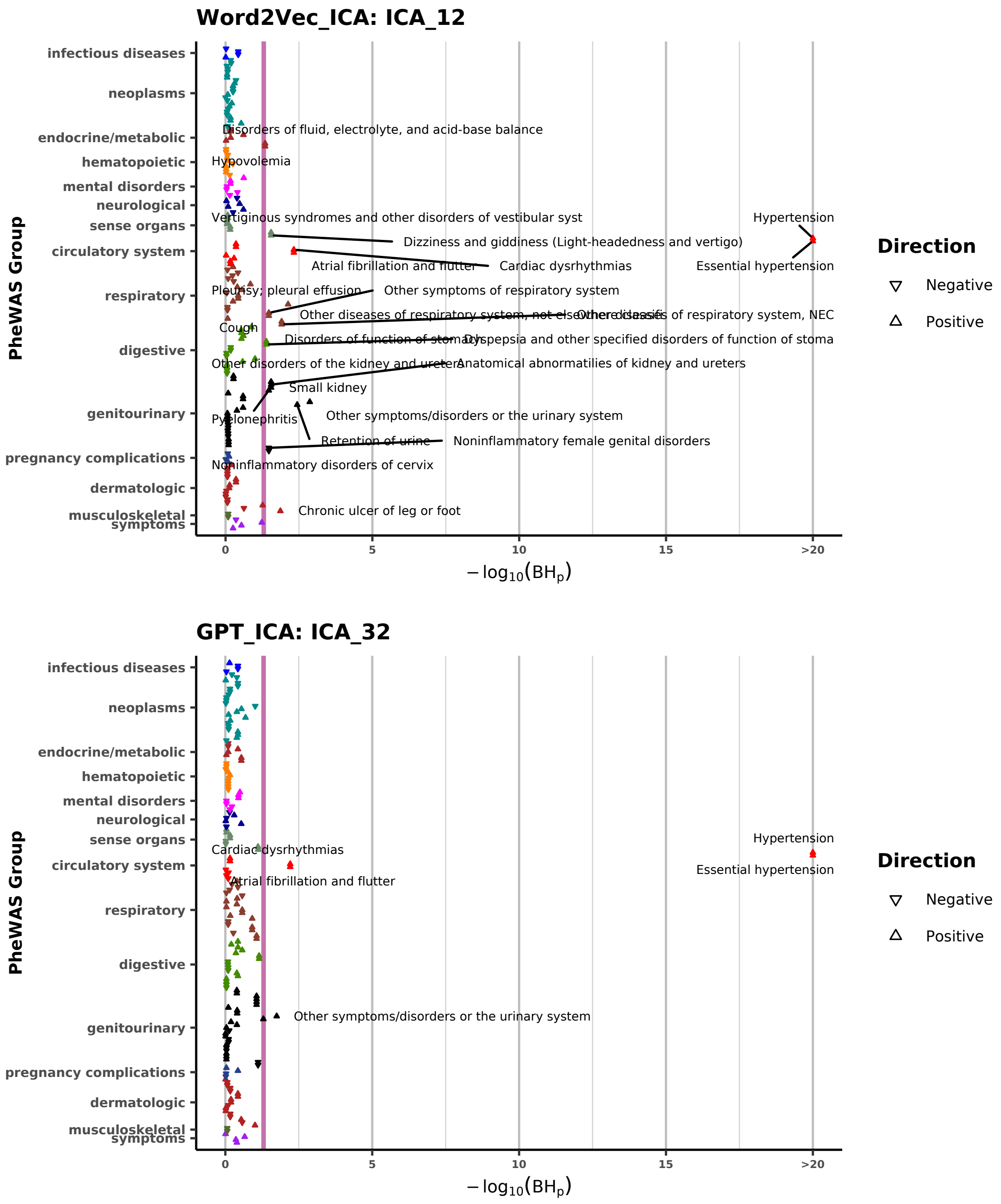

**Supplementary Figure 10. Improved prediction performance of MTAG_EEPRS versus MTAG_PRS.** Bar plots illustrating relative improvements in prediction accuracy of MTAG_EEPRS compared to MTAG_PRS, evaluated using UKBB testing data via 4-fold cross-validation for 13 continuous and 27 binary traits. Relative improvements are quantified as $(R^2_{\text{MTAG\_EEPRS}} - R^2_{\text{MTAG\_PRS}})/R^2_{\text{MTAG\_PRS}}$ for continuous traits and $(\text{AUC}_{\text{MTAG\_EEPRS}} - \text{AUC}_{\text{MTAG\_PRS}})/(\text{AUC}_{\text{MTAG\_PRS}} - 0.5)$ for binary traits, where $R^2$ and AUC values represent the mean across four folds. Horizontal red bars represent improvements for individual traits, and grey bars indicate decreases. Vertical dashed purple lines indicate average improvements across traits, annotated with average improvement percentages.

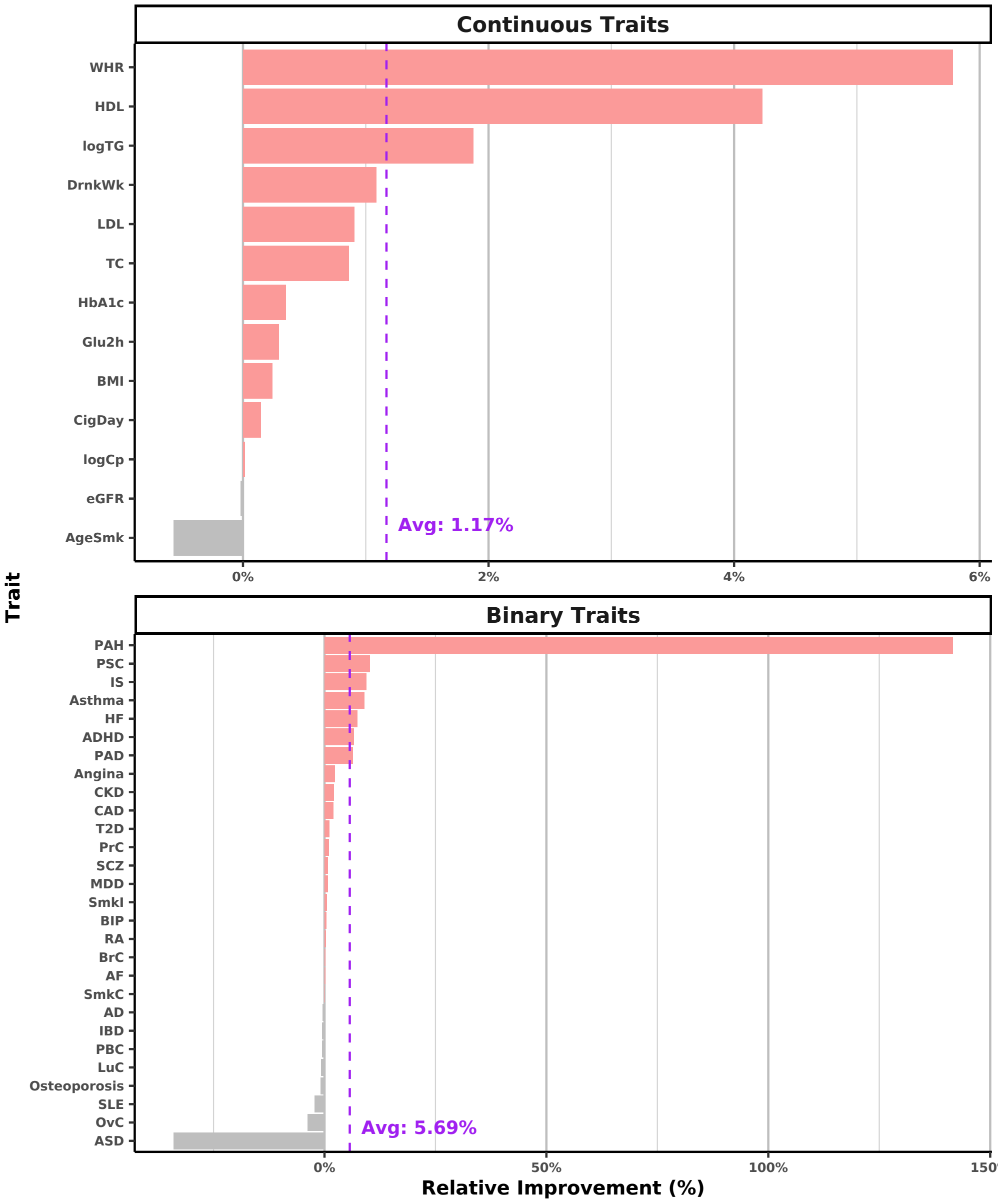
